# The unique topologies of *N^6^*-Adenosine methylation (m^6^A) in land-plant mitochondria and their putative effects on organellar gene-expression

**DOI:** 10.1101/717579

**Authors:** Omer Murik, Sam Aldrin Chandran, Keren Nevo-Dinur, Laure D. Sultan, Corinne Best, Yuval Stein, Carina Hazan, Oren Ostersetzer-Biran

## Abstract

Mitochondria are the main source of ATP production and also contribute to many other processes central to cellular function. Mitochondrial activities have been linked with growth, differentiation and aging. As relicts of bacterial endosymbionts, these organelles contain their own genetic system (i.e., mitogenome or mtDNA). The expression of the mtDNA in plants is complex, particularly at the posttranscriptional level. Following transcription, the polycistronic pre-RNAs in plant mitochondria are processed into individual RNAs, which then undergo extensive modifications, as trimming, splicing and C→U editing, before being translated by organellar ribosomes. Our study focuses on *N^6^*-methylation of Adenosine ribonucleotides (m^6^A-RNA) in plant mitochondria. m^6^A is the most common modification in eukaryotic mRNAs. The biological significance of this highly dynamic modification is under investigation, but it’s widely accepted that m^6^A mediates structural switches that affect RNA stability and activity. By performing m^6^A-pulldown/RNA-seq (m^6^A-RIP-seq) analyses of Arabidopsis and cauliflower mitochondrial transcripts (mtRNAs), we provide with detail information on the m^6^A landscapes in angiosperms mitochondria. The results indicate that m^6^A targets different types of mtRNAs, including coding sequences, UTRs, introns and non-coding RNA species. While introns and noncoding-RNAs undergo multiple m^6^A modifications along the transcript, in mRNAs m^6^A-modifications are preferably positioned near start-codons, and may modulate the translatability of the m^6^A-modified transcripts.

## Introduction

Mitochondria house the oxidative phosphorylation (OXPHOS) system and many metabolic pathways (*e.g*. amino acids, nucleotides and lipids biosynthesis), which are critical to the plant cell (Millar *et al*. 2011, Schertl and Braun 2014). As descendants from an ancestral bacterial symbiont, mitochondria contain their own genome (mtDNA), which encodes a limited set of genes. An exception is the dinoflagellate *Amoebophrya ceratii*, where the entire mitogenome was lost and essential component of the respiratory apparatus were all translocated into the nuclear genome (John *et al*. 2019). These proteins, as well as various organellar cofactors that have been recruited from the host to function in mitochondrial metabolism and organellar gene-expression, are imported into the mitochondria from the cytosol post-translationally (reviewed by e.g., (Fernie *et al*. 2004, Millar, *et al*. 2011, Moller 2001).

The mtDNAs in plants are large and highly variable in size (reviewed by e.g., (Grewe *et al*. 2014, Gualberto *et al*. 2014, Knoop 2012, Skippington *et al*. 2015, Sloan *et al*. 2012, Small 2013). Yet, the number of mitochondrial genes seems more conserved in the land-plant kingdom, with 60∼70 known genes found in different terrestrial plant species (i.e., angiosperms, gymnosperms, ferns, lycophytes, hornworts, mosses and liverworts) (Bonen 2018, Grewe, *et al*. 2014, Gualberto and Newton 2017, Guo *et al*. 2016, Mower *et al*. 2012, Park *et al*. 2015, Sloan, *et al*. 2012). These genes typically include various tRNAs, rRNAs, ribosomal proteins, subunits of the respiratory complexes I (NADH dehydrogenase), III (cytochrome c reductase or bc1), and IV (cytochrome c oxidase), subunits of the ATP-synthase (also denoted as CV), cytochrome *c* biogenesis (CCM) factors, and at least one component of the twin-arginine protein translocation complex (for a recent review see (Bonen 2018). In mistletoe, the mitochondrial genome undergone substantial gene losses including all CI subunits (Maclean *et al*. 2018, Petersen *et al*. 2015, Senkler *et al*. 2018).

The coordination of growth and development in plants is achieved by cellular signaling, which allow them to regulate and coordinate the energy demands during particular growth and developmental stages. These involve both anterograde (nucleus to organelles) and retrograde (organelles to nucleus) signaling cascades (Woodson and Chory 2008), but the identity of the messenger molecules involved in these pathways remains elusive. The expression of mtDNAs in plants is complex, particularly at the posttranscriptional level (reviewed by *e.g*. (Hammani and Giege 2014, Zmudjak and Ostersetzer-Biran 2017). While in Animalia, the mtDNAs are symmetrically transcribed in opposite directions, yielding two large polycistronic RNAs, the mitochondrial genes in plants are arranged in numerous shorted transcription units. Following transcription, the mtRNAs in plants undergo extensive processing steps, which include the removal of many group II-type intron sequences (i.e., splicing) and the conversion of numerous cytidines (C) to uridines (U) deaminations (i.e.,, C→U RNA editing) (reviewed by e.g., (Binder *et al*. 2016, Gualberto and Newton 2017, Hammani and Giege 2014, Liere *et al*. 2011, Small 2013, Zmudjak and Ostersetzer-Biran 2017). These processing steps are critical for the mtRNAs to carry out their functions in protein synthesis, but may also serve as key control points in plant mitochondrial gene-expression, i.e., linking organellar functions with environmental and/or developmental signals.

The manuscript focuses on *N^6^*-adenosine methylation (m^6^A) patterns in the mitochondria of two key Brassicales species, *Arabidopsis thaliana* and *Brassica oleracea* (cauliflower). Various data suggest that m^6^A is the most prevalent RNA modification within mRNA transcripts in eukaryotes (Burgess *et al*. 2016, Liu *et al*. 2015, Niu *et al*. 2013, Patil *et al*. 2018, Visvanathan and Somasundaram 2018), which its biological significance is currently under intense investigation. In animals, m^6^A is a dynamic RNA-base modification which plays important roles for RNA, *e.g*. splicing, stability, and translation, akin to those of DNA methylation and histone modification (Liu, *et al*. 2015, Meyer and Jaffrey 2017, Patil, *et al*. 2018, Visvanathan and Somasundaram 2018). Co-immunoprecipitation of m^6^A-mofifeied transcripts followed by RNA-seq analyses (*i.e*., m^6^A-RIP-seq) provided with detailed information regarding to the m^6^A landscapes in mammalian cells (see *e.g.*, (Dominissini *et al*. 2012). Specific methyltransferases (MTAs, designated as ‘Writers’) and demethylases (termed as ‘Erasers’) regulate the distribution of m^6^A on the transcriptome, whereas proteins that interact and mediate m^6^A-dependent functions are termed as Readers (Meyer and Jaffrey 2017, Patil, *et al*. 2018, Visvanathan and Somasundaram 2018).

In plants, *N^6^*-methylation of Adenosine residues in nuclear-encoded mRNAs is positioned predominantly towards the 3’-end, about 100 bps upstream to the poly A tail (Bodi *et al*. 2012, Burgess, *et al*. 2016, Dominissini, *et al*. 2012, Ke *et al*. 2015, Li *et al*. 2014, Luo *et al*. 2014, Wan *et al*. 2015). This was further supported by a recent RNA-seq data of Arabidopsis plants, using nanopore direct RNA sequencing (Parker *et al*. 2019). Mutant plants with reduced levels of m^6^A, due to the knockout or knockdown of genes homologs of writers and readers show embryogenesis defects and altered growth and developmental phenotypes (Bodi, *et al*. 2012, Shen *et al*. 2016, Zhong *et al*. 2008). Interestingly, global m^6^A-RIP-seq analyses of plant transcriptomes suggested the presence of m^6^A methylations also in plant organelles (i.e., plastid and mitochondrial) as well (Luo, *et al*. 2014, Wan, *et al*. 2015, Wang *et al*. 2017). However, the occurrence and significance of this modification within organellar RNAs are still elusive.

In order to gain detailed insights into the biological significance of this modification in plant organelles, we analyzed the m^6^A patterns of mtRNAs extracted from purified mitochondria obtained from *Arabidopsis thaliana* and *Brassica oleracea* var. botrytis (cauliflower), two key Brassicales species (Grewe, *et al*. 2014). Here, we present several lines of evidence indicating that mtRNAs of Arabidopsis and cauliflower undergo extensive *N^6^*-methyladenosine modifications, and that the m^6^A-RNA methylation affects plant mitochondria gene-expression. Mass-spectroscopy (*i.e*. LC-UV-MS/MS) and m^6^A-RIP-seq analyses indicate that in mitochondria of Arabidopsis and cauliflower plants, the m^6^A is found within different RNA types (*i.e.*, mRNAs, structural RNAs, introns, UTRs and intergenic-expressed regions), but seems particularly prevalent in the noncoding-gene regions. Moreover, while noncoding RNAs contain multiple m^6^A sites found along the transcripts, in mRNAs the m^6^A is predominantly positioned within the AUG translation initiation codon. Biochemical analyses indicate that the extent of m^6^A modifications within the transcript is detrimental for translation. These data strongly support the notion that m^6^A modifications play important roles in regulating the expression of organellar genes in plants.

## Results

### m^6^A is presented within various mitochondrial transcripts in *Arabidopsis thaliana* and *Brassica oleracea* plants

Global RNA-seq analyses revealed the presence of numerous m^6^A modifications within various nuclear-encoded mRNAs in the model plant *Arabidopsis thaliana*, but also suggested that the organellar (*i.e*. chloroplasts and mitochondrial) transcripts undergo extensive *N^6^*-methylations (Bodi, *et al*. 2012, Li, *et al*. 2014, Luo, *et al*. 2014, Shen, *et al*. 2016, Wang, *et al*. 2017). However, the topology of m^6^A in plant mitochondria and significance to organellar biogenesis need to be further investigated. For this purpose, pull-downs of m^6^A-RNAs from highly-enriched mitochondria preparations, obtained from two key Brassicales species (i.e., *Arabidopsis thaliana* seedlings and *Brassica oleracea* (cauliflower) inflorescences), to analyzed by chromatography-mass spectrometry (LC-MS) and RNA-seq analyses. Arabidopsis serves a prime model system for molecular genetics, while cauliflower is employed for the biochemical analysis of plant mitochondria RNA metabolism (Grewe, *et al*. 2014, Keren *et al*. 2009, Neuwirt *et al*. 2005, Sultan *et al*. 2016).

The presence of *N^6^*-methylated Adenosine residues in mitochondrial transcripts (mtRNAs) of Arabidopsis (Luo, *et al*. 2014, Wang, *et al*. 2017) and cauliflower was supported by northern blot analyses (Keren *et al*. 2011) (Supporting information Fig. S1 and %Table S1). Measurements of the bulk levels of m^6^A in total RNA preparations from isolated mitochondria, using a calorimetric m^6^A-RNA quantification kit, indicated to m^6^A/A ratios of 0.44% and 0.38% (*i.e*. 4∼5 m^6^A modifications per 1,000 adenosine nucleotides) (Table S2). These data are in agreement with the reports by Luo, *et al*. (2014) and Wang, *et al*. (2017), which proposed that in global m^6^A-RIP-seq analyses of Arabidopsis the extent of m^6^A modifications within organelles (i.e., chloroplasts and mitochondria) are greater than that of nuclear-encoded mRNAs. Analyses of the relative m^6^A levels, using calorimetric measurements, were previously shown to be well correlated with those of LC-MS analyses (Engel *et al*. 2017, Slobodin *et al*. 2017). Although the use of m^6^A antibodies provide with a sensitive approach to identifying m^6^A methylations, these methods can also lead to erroneous data due to non-specific association of various proteins with the antibodies. LC-MS analyses were used to confirm the presence of m^6^A organellar RNAs isolated from cauliflower mitochondria (Figure 1) (Keren, *et al*. 2009, Neuwirt, *et al*. 2005, Sultan, *et al*. 2016). RNA isolated from purified mitochondria was digested to single nucleosides and the presence of modified Adenosine residues (m^6^A) was assayed by LC-MS, based on specific retention times from standard and modified nucleosides separated by the same chromatographic procedure (see Materials and Methods). As indicated in Figure 2 and supplementary Figure S2, the LC-MS data indicated the presence of *N^6^*-methyladenosine modifications within the transcripts of cauliflower mitochondria.

**Figure 1.**
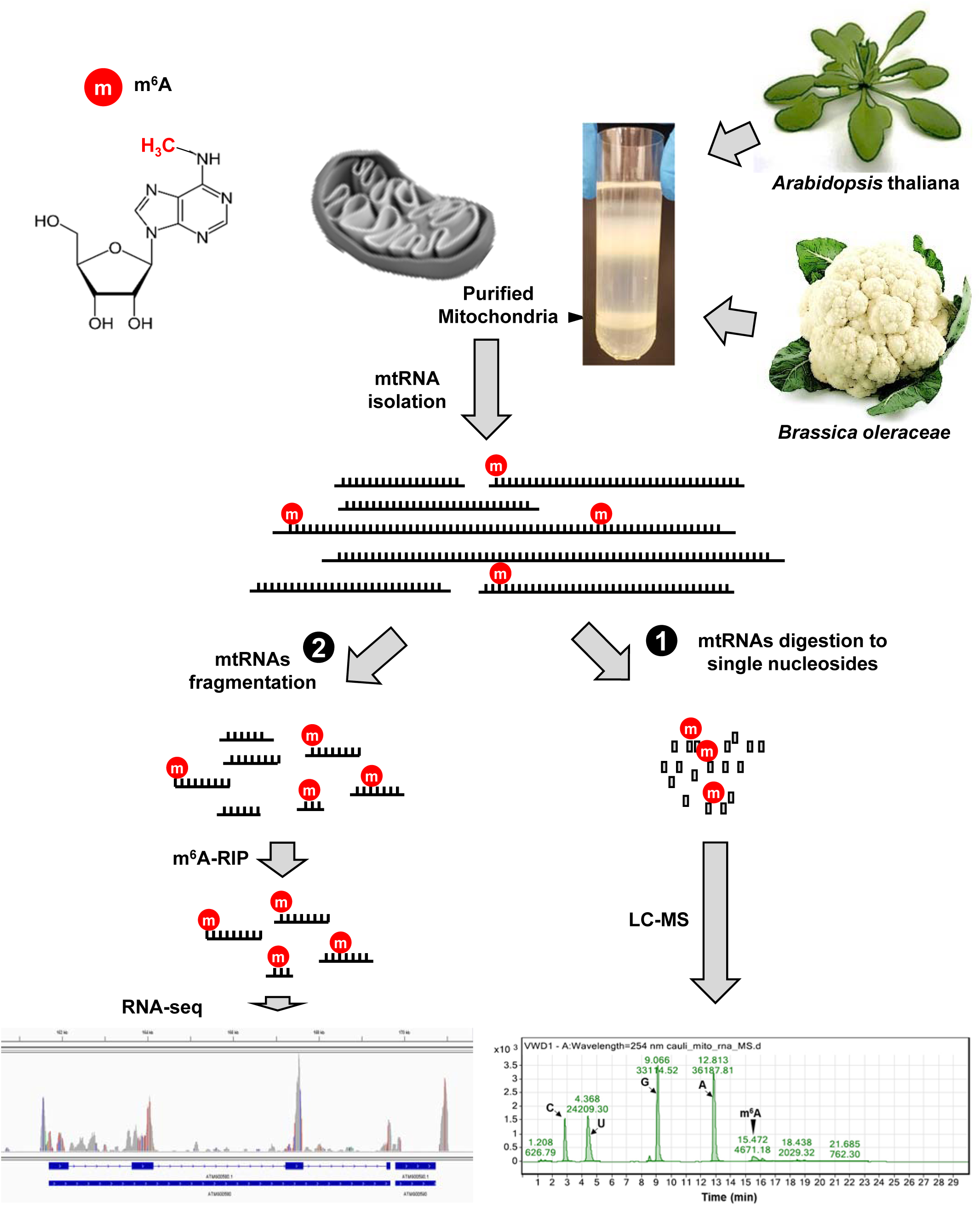
m^6^A-RIP-seq analyses of plant mitochondrial RNAs. Schematic diagram of the m^6^A-seq protocol. Total RNA was isolated from mitochondria obtained from *Arabidopsis thaliana* or Brassica oleracea (var. botrytis; Cauliflower) inflorescences. (1) The presence of *N^6^*-adenosinemethylated (m^6^A) RNAs in the mitochondrial transcripts was confirmed by Liquid chromatography/mass spectrometry (LC-MS) analysis. (2) rRNA-depleted mtRNA was fragmented, and the organellar m^6^A-RNAs were immunoprecipitated, using monoclonal m^6^A antibodies and analyzed by RNA-seq.

**Figure 2.**
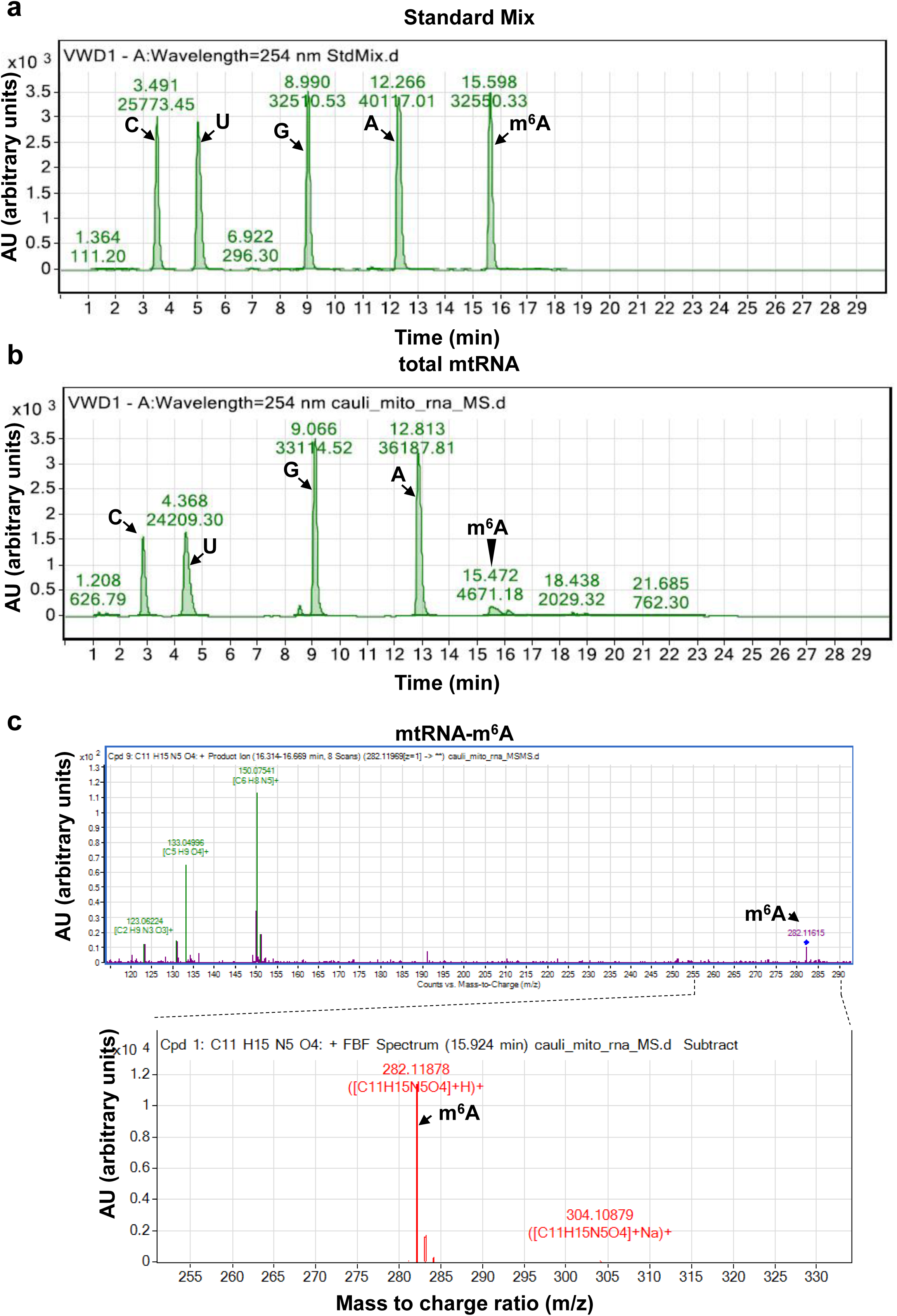
Identification of m^6^A by LC-MS/MS in mitochondrial RNAs isolated from wild-type cauliflower inflorescences. Total mtRNA, isolated from cauliflower mitochondria, was digested to single nucleosides with S1 nuclease and phosphodiesterase, and the presence of modified Adenosine residues (m^6^A) was assayed by LC-MS, based on specific retention times from standard and modified nucleosides separated by the same chromatographic procedure. The UV absorption spectra of (a) different nucleosides (A, G, C, and U) and m^6^A standards, (b) nucleosides obtained by total RNA digestion and (c) the specific MS spectrum of m^6^A in mitochondrial RNA preps are given in each panel. The LC-MS total ion chromatogram data of mtRNA nucleosides is presented in Figure S2. The chromatographic data, including the retention times (RT), masses, abundances and molecular formula of the analyzed products are summarized within Table S1

**Table 1.** m^6^A-RIP-seq data

**(a)** m<sup>6</sup>A-RIP-seq data corresponding to SRA accessions PRJNA472433, replicate 1.
| m<sup>6</sup>A-RNA |  | *Arabidopsis thaliana*<br>(No. of peaks) | *Brassica oleracea*<br>(No. of peaks) |
| --- | --- | --- | --- |
| Known genes |  | 189 peaks / 41 genes | 310 peaks / 18 genes |
|  | UTRs | 8 | 17 |
|  | Exons/coding regions | 115 | 162 |
|  | Introns | 66 | 131 |
| mtORFs |  | 11 peaks / 10 genes | 51 peaks / 21 genes |
| Intergenic regions |  | 195 | 349 |
| rRNAs*<sup>1</sup> |  | (12) | (9) |
| tRNAs*<sup>1</sup> |  | (1) | (8) |

**(b)** m<sup>6</sup>A-RIP-seq data corresponding to SRA accessions PRJNA472433, replicate 2.
| m<sup>6</sup>A-RNA |  | *Arabidopsis thaliana*<br>(No. of peaks) | *Brassica oleracea*<br>(No. of peaks) |
| --- | --- | --- | --- |
| Known genes |  | 235 peaks / 37 genes | 266 peaks / 23 genes |
|  | UTRs | 27 | 12 |
|  | Exons/coding regions | 122 | 150 |
|  | Introns | 85 | 104 |
| mtORFs |  | 16 peaks / 7 genes | 35 peaks / 22 genes |
| Intergenic regions |  | 244 | 305 |
| rRNAs*<sup>1</sup> |  | (18) | (2) |
| tRNAs*<sup>1</sup> |  | (1) | (6) |

\*<sup>1</sup> - The m<sup>6</sup>A-RIP-seq data corresponding to rRNAs and tRNAs should be taken with care due to rRNA and tRNA depletion prior to the RNA-seq analyses (see Material and Methods).

### The unique topologies of m^6^A methylomes in Brassicales mitochondria

Following the confirmation of *N^6^*-adenosine methylation events (Fig. 2, and supplementary Figures S1 and S2), we further characterized the landscapes of m^6^A modifications in Arabidopsis and cauliflower mitochondria by the m^6^A-RIP-seq method (Dominissini, *et al*. 2012, Luo, *et al*. 2014) (Fig. 1). rRNA-depleted RNA preparations, obtained from Arabidopsis and cauliflower mitochondria, were fragmented into 60∼200 nucleotides (*i.e*. input mtRNA) and immunoprecipitated with monoclonal anti-m^6^A antibodies (see Materials and Methods). The precipitated m^6^A-mtRNAs were then used to construct libraries that were subjected to RNA-seq analyses. The corresponding nucleotide reads were aligned to the reference Arabidopsis and cauliflower mitochondrial transcriptomes (Fig. S3) (Grewe, *et al*. 2014, Sultan, *et al*. 2016, Unseld *et al*. 1997) to generate detailed landscapes of m^6^A-RNAs in Arabidopsis and cauliflower mitochondria. As control we analyzed the RNA-seq’s of rRNA-depleted mtRNAs from Arabidopsis and cauliflower (SRA accession PRJNA472433, Table S3).

The mitogenome of *Arabidopsis thaliana* (Col-0) harbors 57 identified genes, including 38 protein coding frames, 22 tRNAs and 3 rRNAs (Sloan *et al*. 2018, Unseld, *et al*. 1997). The remaining four genes include an intronic ORF, annotated as maturase-related (*matR*) gene encoded in the fourth intron in *nad1* (Sultan, *et al*. 2016, Wahleithner *et al*. 1990), *atp4* (previously annotated as *orf25*) (Heazlewood *et al*. 2003), *atp8* (previously denoted as *orfB*) (Heazlewood, *et al*. 2003, Sabar *et al*. 2003) and *mttB* (or *tatC*, previously annotated as orfX) (Bogsch *et al*. 1998). In addition to known genes, about 460 open reading frames (i.e., mtORFs; larger than 60 codons) are found within the sequenced mtDNA of *Arabidopsis thaliana* (Col-0) (Unseld, *et al*. 1997). The m^6^A/RNA-seq data is presented Table 1 and supplementary Tables S3 and S4, which corresponds to two independent m^6^A-RIP-seq analyses. The m^6^A modifications were mapped to 37 ∼ 41 annotated genes and 7 ∼ 10 ORFs in the mitogenome of Arabidopsis (Table 1 and Tables S3 and S4).

In cauliflower mitochondria, 54 known genes (i.e., 33 annotated protein coding frames, 18 tRNAs and 3 rRNAs) and 35 ORFs are annotated (KJ820683.1) (Grewe, *et al*. 2014). Between 18 to 23 genes and 21 or 22 mtORFs were identified in the two replicates of the m^6^A-RIP-seq analyses of *B. oleracea* mitochondria (see Table 1, and supplementary Tables S3 and S4). The distribution of m^6^A signals in Arabidopsis and cauliflower mitochondria (i.e., average of the two independent m^6^A-RIP-seq analyses) is provided in Figure 3.

**Figure 3.**
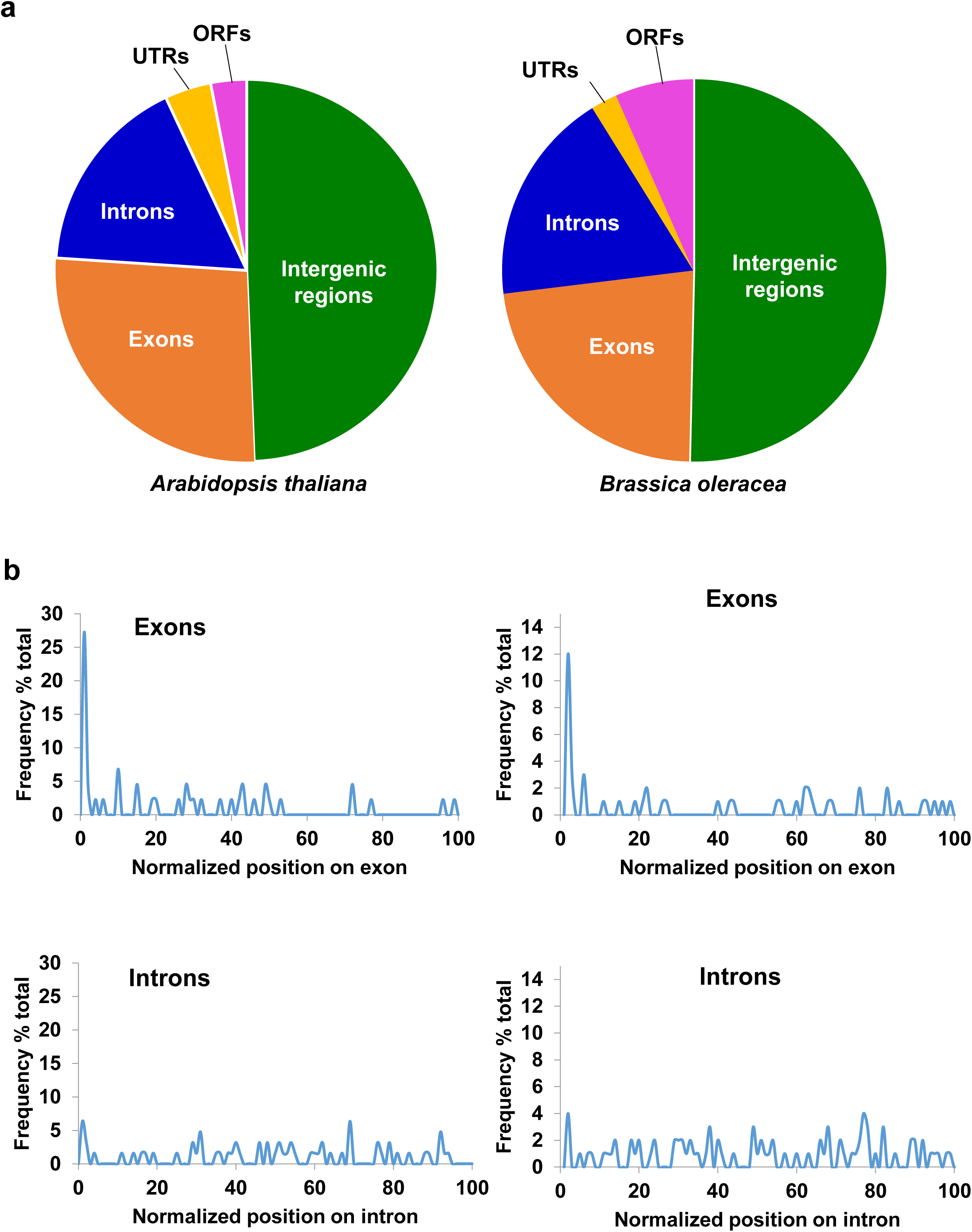
The m^6^A landscapes of plant mtRNAs reveal a unique topology. (a) pie-charts presenting the fraction of m^6^A peaks in coding-regions (exons), introns, UTRs and intergenic regions in Arabidopsis (left panel) or Cauliflower (right panel) mitochondria, represented in the m6A-RNA-seq data (*i.e.*, SRA accessions PRJNA472433, replicate 1). (b) Distribution of m^6^A peaks in Arabidopsis (left panel) or Cauliflower (right panel) sequences corresponding to exons (upper panels) and intron regions (lower panels), according to normalized exons and introns lengths (PRJNA472433, replicate 1). A list of mitochondrial genes in Arabidopsis and cauliflower identified by the m^6^A-RIP-seq analyses is presented in Table S4.

**Figure 4.**
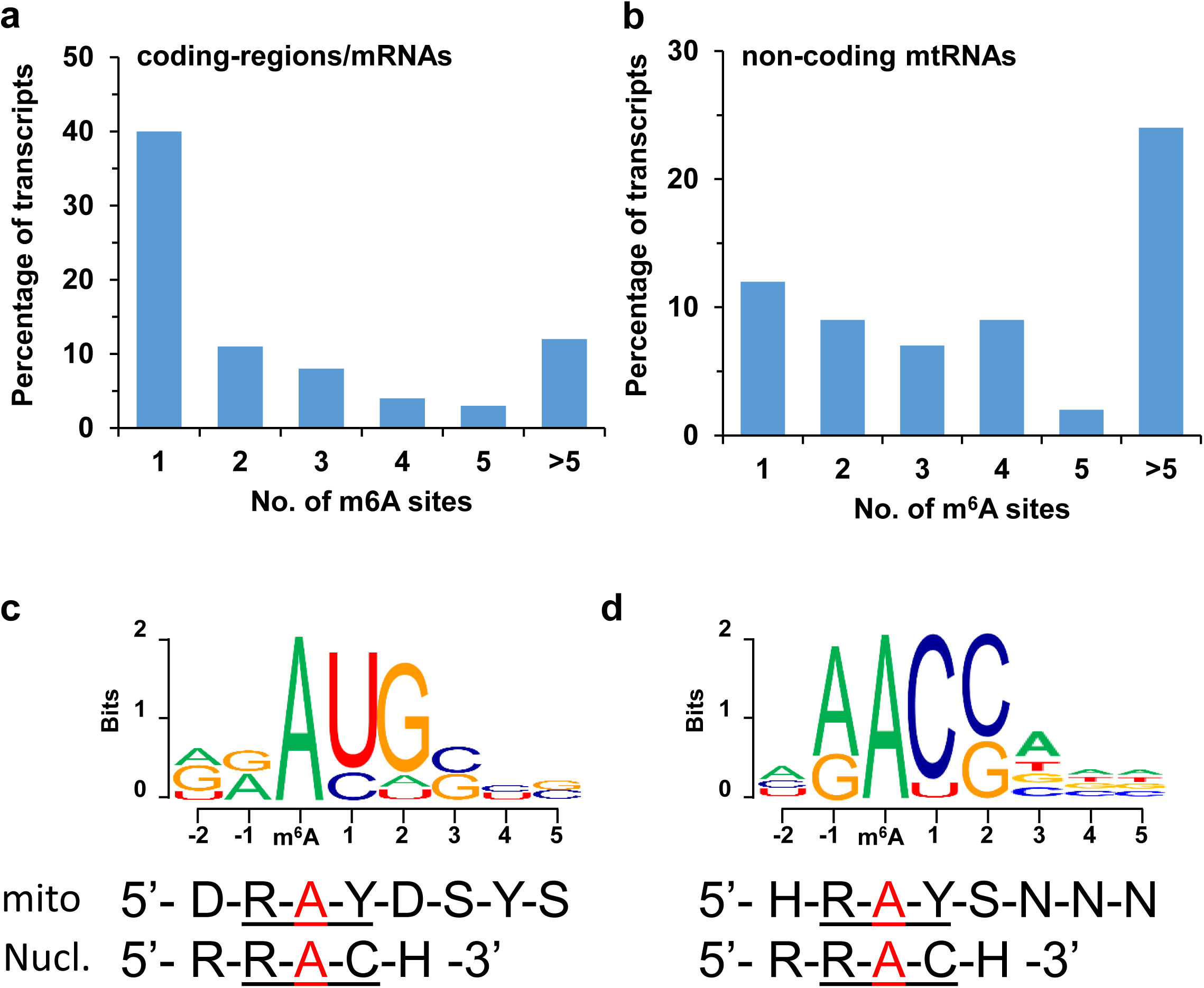
Common m^6^A patterns in Arabidopsis and cauliflower mitochondria. Relative abundances of *N^6^*-adenosinemethylated nucleotide sites in organellar transcripts in both Arabidopsis and cauliflower, corresponding to (a) coding-regions (or exons) and (b) non-coding RNAs. The identity of putative consensus sequence motifs in m^6^A-modified mtRNA, was analyzed by the HOMER (Heinz, *et al*. 2010) and MEME (Bailey, *et al*. 2009) programs. Integration of results from the motif discovery tools, indicated to the common m^6^A elements, DRm^6^AYHSYS (where ‘D’ is A, G or U, ‘R’ is A or G, ‘Y’ is C or U, ‘S’ is C or G, and ‘H’ is A, C or U) for mRNAs (c) and HRm^6^AYS for noncoding mtRNAs (d).

The identity of the genes and the numbers of modifications within each gene were different between Arabidopsis and cauliflower, but the m^6^A-related patterns in Arabidopsis and cauliflower mitochondria generally shared similar patterns. In animals, m^6^A sites are in particularly enriched within the 3’ UTRs, near stop codons (Bodi, *et al*. 2012, Deng *et al*. 2015, Dominissini, *et al*. 2012, Hoernes *et al*. 2016). Yet, in both Arabidopsis and cauliflower, the majority of the m^6^A peaks (i.e., about half of the signals) are mainly found in noncoding RNAs (49.3 ± 7.8 % in Arabidopsis and 50.3 ± 4.8 % in cauliflower) (Fig. 3a, Table 1 and supplementary Table S4). Within known gene regions, about quarter to fifth of the m^6^A methylation sites are mapped to the coding sequences (i.e., 26.6 ± 1.1 % in Arabidopsis and 22.7 ± 0.5 % in cauliflower), about 20% are found within group II intron sequences (17.0 ± 3.0 % in Arabidopsis and 18.1 ± 2.9 % in cauliflower), and only 2 to 4 % (3.9 ± 3.0 % in Arabidopsis and 2.2 ± 0.5 % in cauliflower) of the sites occur within the untranslated regions (UTRs, Fig. 3a). Higher m^6^A signals in mRNAs, versus introns and other non-coding regions (Fig. 3 and Fig. S3), may be expected from their expression patterns and RNA metabolism in plants mitochondria (Grewe, *et al*. 2014, Zmudjak and Ostersetzer-Biran 2017). As these analyses were performed with rRNA-depleted mtRNAs, any conclusions regarding the presence of m^6^A in tRNAs and rRNAs (Table 1) should be taken with care. The differences in the m^6^A landscapes between the two closely-related Brassicales species may correspond to variations in the experimental procedure, or indicate to altered m^6^A modifications due to growth conditions and the use of different plant tissues (i.e., vegetative plant tissue in Arabidopsis and floral tissues in cauliflower). Analyses of the m^6^A landscapes in different tissues and at various growth stages are underway.

To evaluate the m^6^A-enrichment within coding sequences systematically, we aligned the m^6^A peaks of Arabidopsis and cauliflower mitochondria to their mtDNA sequences. These included coding sequences, group II introns and noncoding regions. While in introns (Fig. 3b) and intergenic regions (Fig. S4) the m^6^A signals are distributed along the transcript, in protein coding sequences (i.e., exons and mRNAs) the m^6^A sites are predominantly located within the range of ±100 nucleotides around the first codon (i.e., about 25% of the reads in Arabidopsis and 12% in cauliflower mitochondria) (Fig. 3a,b). These data may indicate that in mRNAs of both Arabidopsis and cauliflower mitochondria, many of the m^6^A modifications occur in proximity or within the translation initiation site. We also noted to differences in the quantities of m^6^A modifications per organellar transcript between mRNAs and noncoding transcripts. Many mRNAs contain a single or a few m^6^A modification site (Fig. 3b and Fig. 4a), while in noncoding sequences often contain more than 5 sites per transcript (see Fig. 3b and Fig. 4b). In summary, the m^6^A methylation patterns in plant mitochondria differ markedly from those of bacteria, where the m^6^A is found mostly inside ORFs, or nuclear-encoded mRNAs which are modified mainly in the 3’ UTRs and regions adjacent to the stop codon (Bodi, *et al*. 2012, Deng, *et al*. 2015, Dominissini, *et al*. 2012, Hoernes, *et al*. 2016).

### Analysis of common m^6^A motifs in Brassicales mitochondrial RNAs

To determine whether the m^6^A-RNA reads in plant mitochondria share a common sequence motif, we used the HOMER (Heinz *et al*. 2010) and MEME (Bailey *et al*. 2009) programs. Integration of results from the motif discovery tools, and clustering of the enriched mRNAs or noncoding sequences from Arabidopsis and cauliflower mitochondria, suggested the following m^6^A elements: DR<u>m^6^A</u>YHSYS (where ‘D’ is A, G or U, ‘R’ is A or G, ‘Y’ is C or U, ‘S’ is C or G, and ‘H’ is A, C or U) for mRNAs (Fig. 4c) and HR<u>m^6^A</u>YS for the noncoding RNAs (Fig. 4d). These putative mitochondrial <u>m^6^A</u> sequence elements share a similar core motif, *i.e*., R<u>m^6^A</u>Y (Fig. 4c,d), which was described for nuclear-encoded mRNAs both animals and plants (*i.e*., RR<u>m^6^A</u>CH) in (Liu and Pan 2016, Maity and Das 2016, Patil, *et al*. 2018), and to a lesser degree with the <u>m^6^A</u> motif (*i.e*., UGCC<u>m^6^A</u>G) of bacterial RNAs (Deng, *et al*. 2015).

The consensus DR<u>m^6^A</u>YHSYS and HR<u>m^6^A</u>YS sequences are somewhat different from the previously suggested <u>m^6^A</u> motif (i.e., RR<u>m^6^A</u>CH site) for organelles in Arabidopsis plants (Wang, *et al*. 2017). The variations between these analyses may relate to the use of <u>m^6^A</u>-RNA-seq with global RNA-seq analyses in Wang, *et al*. (2017), which might be only partially represented by mitochondrial transcripts.

### The m^6^A modification has no obvious effects on the stability of recombinant transcripts, in vitro

The identity of the m^6^A-modifying enzymes in bacteria or organelles in eukaryotic organisms is currently unknown. To gain some insights regarding the physiological significance of m^6^A in angiosperms mitochondria, we analyzed the effects of *N^6^*-adenosine methylations on the transcription, stability and translatability of several mtRNAs, in vitro. For this purpose, cDNA fragments corresponding to ATP-synthase *atp1* gene, *nad4* subunit of respiratory complex I (i.e., NADH dehydrogenase enzyme), whose coding region is interrupted by several group II-type introns in both Arabidopsis and cauliflower mitochondria (Grewe, *et al*. 2014, Sloan, *et al*. 2018, Unseld, *et al*. 1997), and *matR* ORF, encoded within *nad1* intron 4 (Sultan, *et al*. 2016, Wahleithner, *et al*. 1990), were obtained by RT-PCR with total RNA isolated from Arabidopsis mitochondria and cloned into pGEM vector, such as each gene-fragment contains a T7 RNA polymerase (RNAP) promoter site at the 5’ termini and a C-terminal 6x His tag in its 3’ termini (Fig. 5, and Materials and Methods). The use of RT-PCR instead of PCR allowed us to retain potentially important C→U RNA editing sites, which are highly common in plant mitochondria and affect *atp1*, *nad4* and *matR* mRNAs (Hammani and Giege 2014, Ichinose and Sugita 2016b, Sloan, *et al*. 2018, Takenaka *et al*. 2008, Unseld, *et al*. 1997, Zmudjak and Ostersetzer-Biran 2017). The integrity of each construct was verified by DNA-sequencing, and then transcribed in vitro, in the absence or presence of different m^6^ATP/ATP nucleotides ratios, in order to obtain ‘body-labeled’ m^6^A-modified transcripts (Fig. 5).

**Figure 5.**
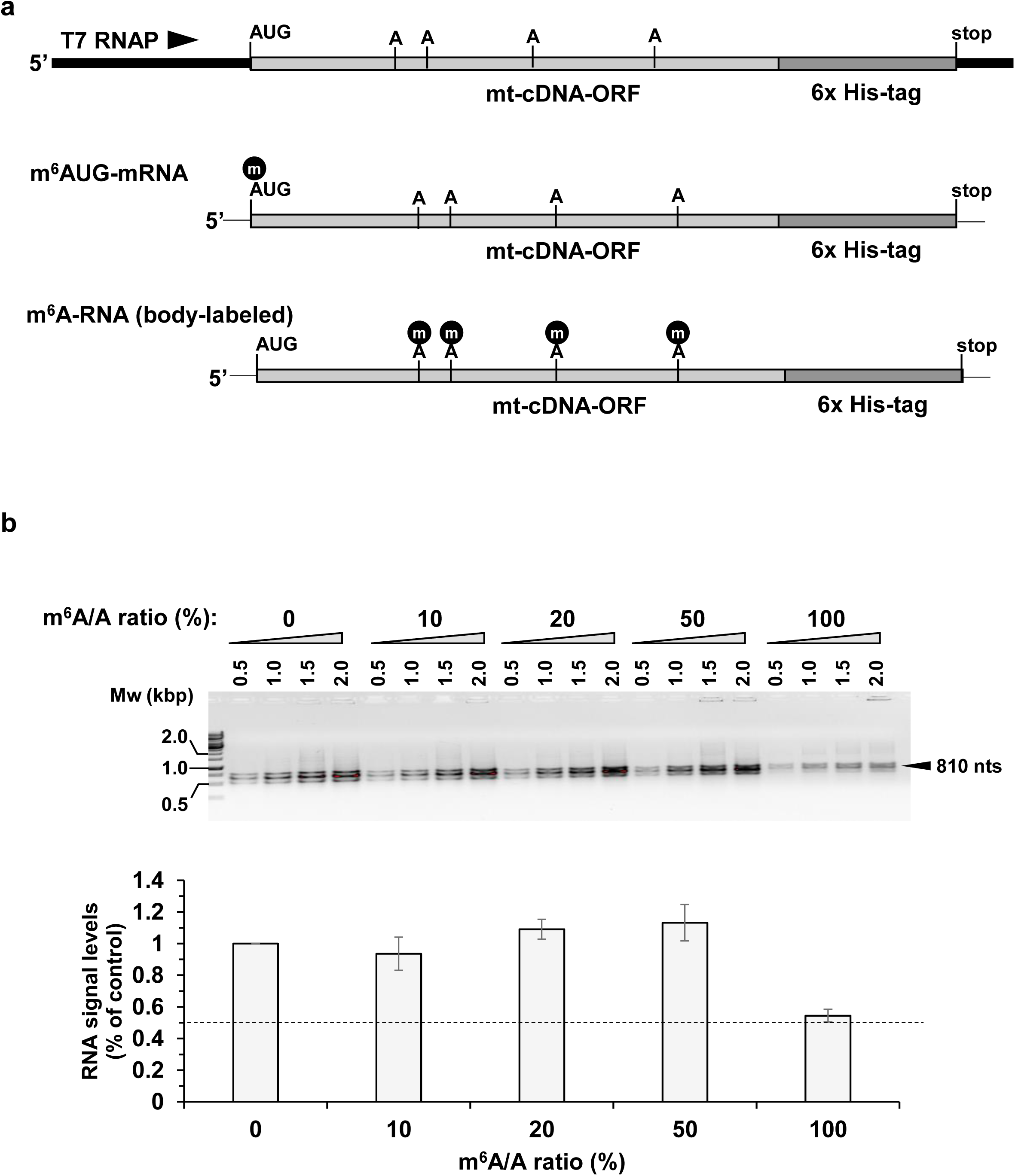
In vitro transcription of body-labeled m^6^A-RNAs. (a) Schematic representation of constructs used for in vitro transcription assays. cDNAs corresponding to *atp1*, *matR* and *nad4* were obtained by RT-PCR, with total RNA isolated from Arabidopsis mitochondria, such as each gene-fragment contained a T7 RNA polymerase (RNAP) promoter site at the 5’ termini and a C-terminal 6x His tag (see Materials and Methods). mtRNAs modified only in the initiation codon (m^6^AUG-mRNA) were generated by custom RNA synthesis (see Materials and Methods). (b) *In vitro* transcription of cDNAs (*i.e*., *atp1*) with T7 RNA polymerase, in the absence (0) or presence of 10, 20, 50 or 100% m^6^ATP/ATP ratios, were used in order to obtain ‘body-labeled’ m^6^A transcripts. Different amounts (*i.e.,* 0.5, 1.0, 1.5 or 2.0 µl) of *atp1* transcripts (diluted to a final RNA conc. of 100 ng/μl) were loaded onto agarose gel (1% w/v), and separated by gel-electrophoresis. Quantification data of the relative levels of *atp1* mRNA transcripts are indicated below the blots in panel ‘b’.

The decay patterns of control (*i.e*., unmodified) versus m^6^A-modified *atp1*, *nad4* and *matR* RNAs were examined in vitro, by incubating the transcripts with purified cauliflower mitochondrial extracts. Samples were collected at different time points, as indicated in Figure 6 and supplementary Figure S5. As indicated by the ‘*in organello*’ assays, the presence of m^6^A had no or only minor effect on the stabilities of *atp1*, *nad4* and *matR* transcripts (Fig. 6 and Fig. S5). Both the control RNAs and the m^6^A-modified transcripts showed similar degradation patterns, where the signal of the recombinant RNAs was reduced by about 40∼50%, following two hours of incubation with the plant orgaenllar extracts (Figs. 6b and S5).

**Figure 6.**
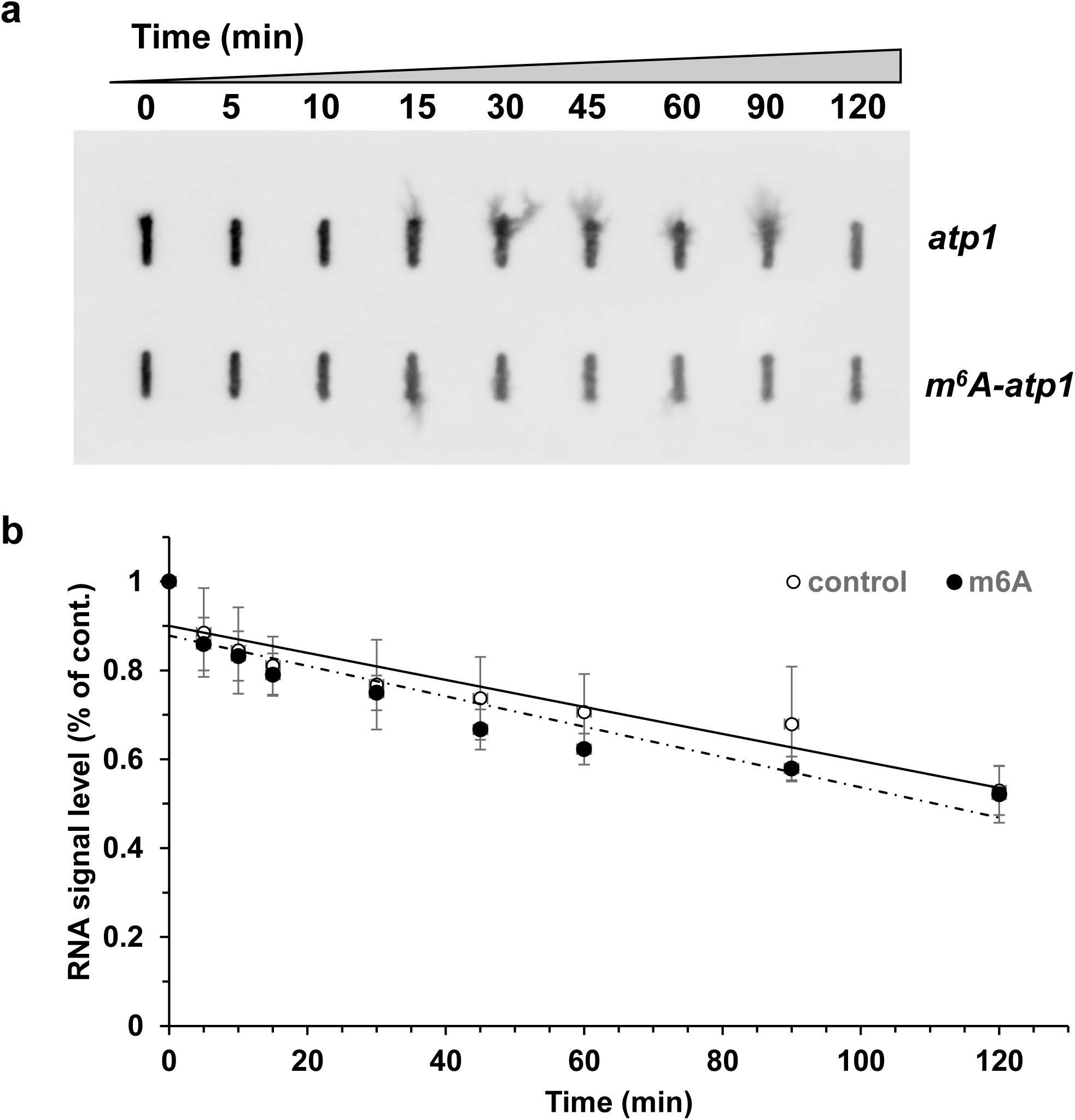
The effect of m^6^A modifications on mtRNA stability. (a) The decay patterns of control (*i.e*., non-modified) versus m^6^A body labeled mRNAs, were examined by incubating the in vitro transcribed *atp1* mRNAs with purified cauliflower mitochondrial extracts. Samples were collected at different time points, as indicated in the figure, and the relative levels of atp1 mRNAs were examined by Filter binding assays (Keren, *et al*. 2008, Keren, *et al*. 2011, Ostersetzer, *et al*. 2005). (b) Quantification data of the relative levels of *atp1* mRNA transcripts following incubation with the organellar extracts. The The RNA-degradation patterns of m^6^A modified versus control (non-modified) *matR* and *nad4* mRNAs are presented in Figure S5.

### m^6^A affects the translation of mitochondrial transcripts, *in vitro*

Next, we considered a possible link between posttranscriptional m^6^A modification and organellar translation (Akichika *et al*. 2019, Choi *et al*. 2016, Hoernes, *et al*. 2016, Meyer and Jaffrey 2017, Meyer *et al*. 2015, Slobodin, *et al*. 2017, Sun *et al*. 2019). Attempts to induce the translation of in vitro transcribed RNAs in Arabidopsis or cauliflower mitochondrial extracts were generally unsuccessful. We therefore analyzed the translation efficiencies of m^6^A-modified *atp1*, *nad4* and *matR* transcripts (Fig. 5), using the TNT^®^ wheat germ extract (WGE) system (see Materials and Methods). To enable the synthesis of recombinant proteins, specifically by organellar ribosomes (i.e., the crude wheat germ extracts should contain both cytosolic and organellar ribosomes), the translation of recombinant ATP1, MATR and NAD4 fragments was assayed in the presence of cycloheximide (CycH, 100 μg/ml), which blocks translation by affecting peptidyl transferases activity in eukaryotic ribosomes, and Kanamycin (Kan, 50 μg/ml) which specifically interrupt organellar (*i.e*., mitochondria and plastids) proteostasis (Wang *et al*. 2015a).

As indicated in Figure 7a, the translation of *atp1*, *matR* and *nad4* was reduced in the presence of CycH by about 50% when the transcripts contained a 10% m^6^A/A ratio, while no translation products corresponding to *atp1*, *matR* and *nad4* were observed when their transcripts were synthesized in the presence of m^6^A only (*i.e*., 100% m^6^A/A ratio). Excessive m^6^A modification was found to be detrimental for the translation of m^6^A modified *atp1*, *matR* and *nad4 in vitro* (*i.e.*, in the WGE system), as indicated by the reduced translation efficiencies of the modified mtRNAs containing a higher m^6^A/A ratio. As indicated in Figure 7a, the addition of Kanamycin abolished the translation by organellar ribosome in the plant extract.

**Figure 7.**
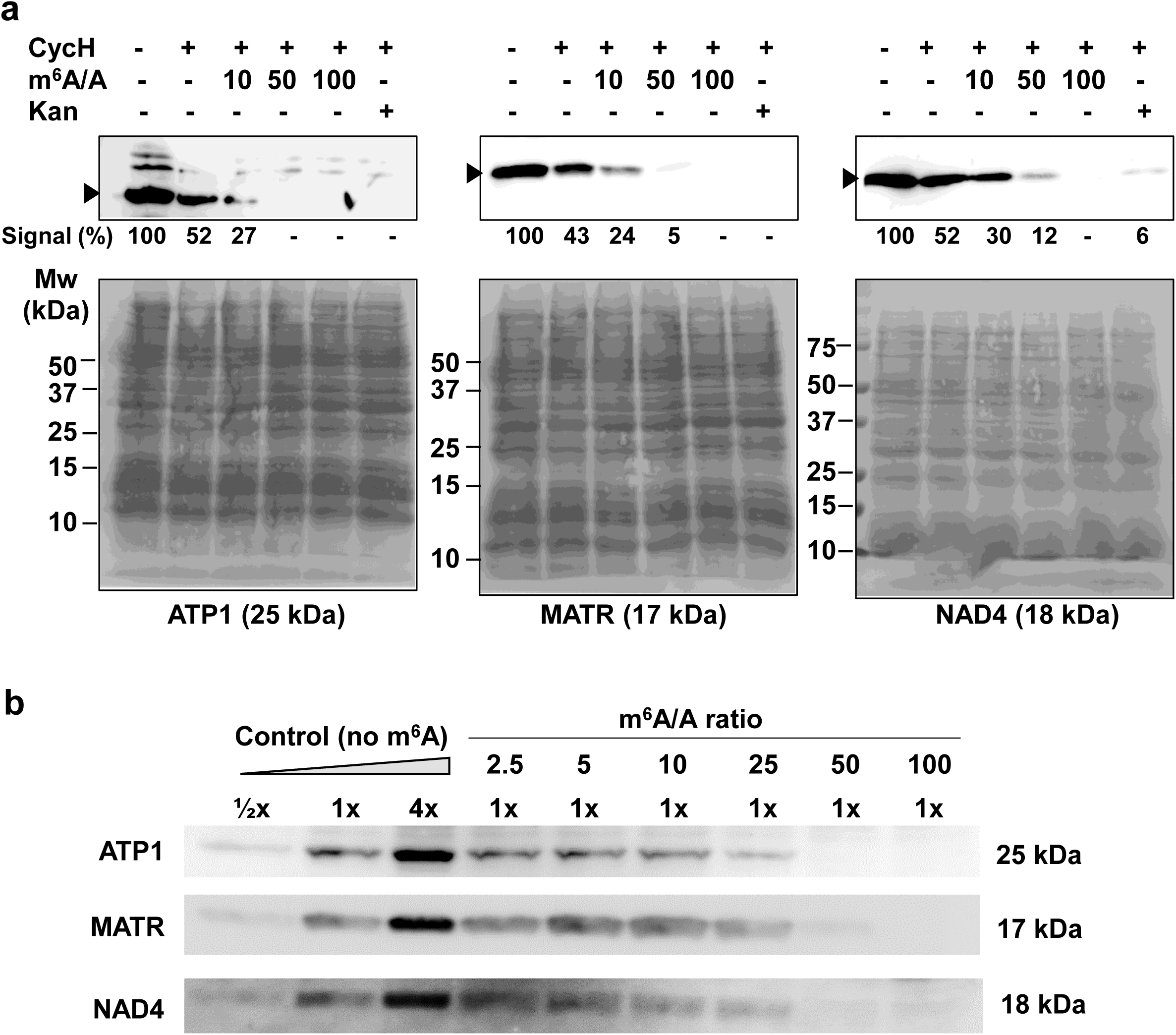
m^6^A is detrimental for the translation of mtRNAs modified at multiple sites. (a) The translation efficiencies of transcripts corresponding to *atp1*, *nad4* and *matR*, containing increased levels of m^6^A modifications along the transcripts (*i.e*., body-labeled RNAs), were examined by western-blot analyses of in vitro translated products, using a wheat germ extract system. To enable the synthesis of recombinant proteins by organellar ribosomes, the translation of the corresponding *atp1*, *matR* and *nad4* transcripts was assayed in the presence of cycloheximide (CycH, 100 μg/ml), which blocks peptidyl transferase activity in eukaryotic ribosomes. In addition, the translation efficiencies were also analyzed in the presence of Kanamycin (50 μg/ml), which interrupt the synthesis of organellar (*i.e.*, mitochondria and plastids) proteins (Wang, *et al*. 2015a). (b) The translation efficiencies of *atp1*, *matR* and *nad4* transcripts, modified in the presence of different, *i.e.*, 0, 2.5, 5, 10, 25, 50 and 100%) m^6^A/A ratios.

We further analyzed the effects of m^6^A on the translation of *atp1*, *matR* and *nad4* transcripts specifically modified at their AUG sites (i.e., m^6^AUG). For this purpose, synthetic RNA fragments, containing the consensus T7 polymerase promoter upstream to a modified Adenosine residue of the imitation codon (i.e., TTTAA<u>GAAGGAG</u>ATATACC[^m6^A]TG-xxxx; xxx indicates the *atp1*, *nad4* or *matR* fragments), were translated in the absence or the presence of cycloheximide (CycH, 100 μg/ml). We noticed that the efficiency of translation of *atp1* and *nad4* transcripts, modified only at the Adenosine residue of the initiation codon, was increased to some extent in the WGE (Fig. 8a). Yet, when the translation efficiencies of the m^6^AUG-RNAs were assayed in the presence of Kanamycin, similar intensities of the His-tagged polypeptides, corresponding to ATP1, MATR and NAD4 fragments, were seen in the non-modified and m^6^A-AUG modified transcripts. These analyses indicate that excessive m^6^A modification is detrimental for the translation of m6A-modified transcripts, while a single modification at the start codon may enhance to some extent the trainability by plant organellar ribosomes.

**Figure 8.**
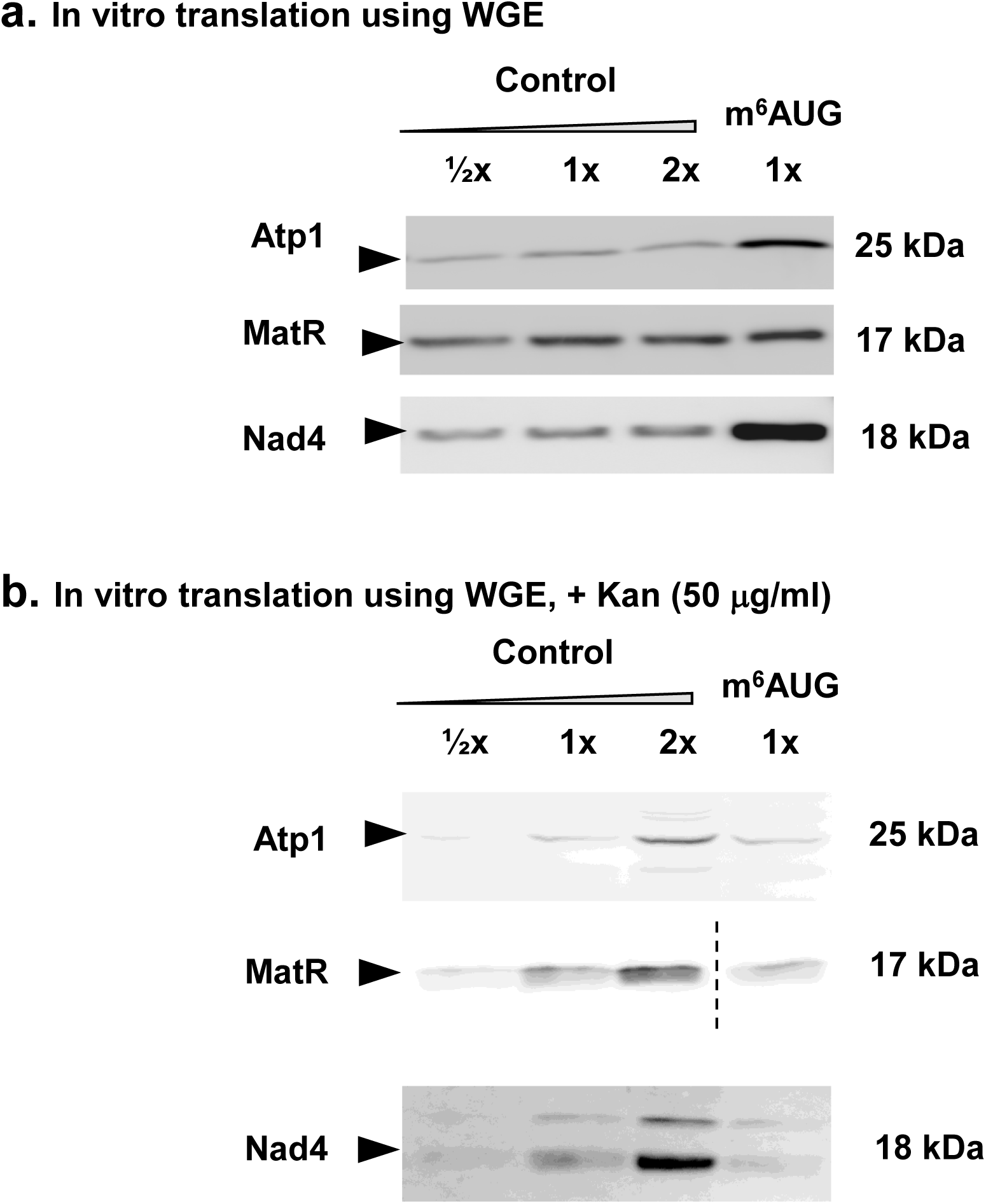
The translation efficiencies of transcripts corresponding to *atp1*, *nad4* and *matR*, modified specifically at the AUG site (*i.e*., m^6^AUG). (a) The translation efficiencies of transcripts corresponding to *atp1*, *nad4* and *matR*, modified by m^6^A specifically at the AUG sites, were examined by western-blot analyses of in vitro translated products, using a wheat germ extract system. (b) To enable the synthesis of recombinant proteins by organellar ribosomes, the translation of the corresponding *atp1*, *matR* and *nad4* transcripts was assayed in the presence of in the presence of Kanamycin (50 μg/ml), which interfere with the synthesis of mitochondrial (or plastidial) polypeptides (Wang, *et al*. 2015a). The authors wish to note that the image in the MATR panel in Fig. 8b was cropped in order to bring the m^6^A-MatR lane close to the control (unmodified) lanes (marked by a dashed-line). No other changes have been made to the original figure.

### Arabidopsis mutants affected in mitochondria gene expression show altered m^6^A patterns

The biogenesis of mitochondria in plants is tightly controlled by nuclear loci, which regulate organellar gene expression and metabolic functions. The plant organeller pre-RNAs undergo extensive posttranscriptional processing steps, including RNA editing and intron splicing. These essential processing steps are regulated by various cofactors (reviewed by *e.g*. (Zmudjak and Ostersetzer-Biran 2017), such as the maturase-related nMAT1 (Keren *et al*. 2012, Nakagawa and Sakurai 2006) and nMAT2 (Keren, *et al*. 2009, Zmudjak *et al*. 2017) proteins, which affect the splicing efficiencies of different subsets of mitochondrial introns in Arabidopsis. Both *nmat1* and *nmat2* mutants show altered mitochondria gene expression profiles, with notable accumulations of various group II intron-containing pre-RNAs and decreased steady-state levels of the corresponding mRNAs (Keren, *et al*. 2009, Keren, *et al*. 2012, Nakagawa and Sakurai 2006, Zmudjak, *et al*. 2017).

m^6^A-RIP analyses followed by RT-qPCRs of *nmat1*, *nmat2* and wild-type plants suggested that pre-RNAs that their maturation (i.e., splicing) was affected are also more extensively modified by m^6^A (Fig. 9 and Table S5). These including the first exons in *nad1* (*i.e*., *nad1-*ab), *nad2-*ab and *nad4-*bc RNAs in *nmat1* mutant, and *cox2*, *nad1-*de, *nad7-*bc, and to a lesser degree the different exons in *nad2* and *nad5* genes, and the intronless transcripts of *ccmB*, *ccmC* and *nad9* mRNAs in *nmat2* mutant (Fig. 9). Higher accumulation was observed in the levels of *nad1*-i1, *nad2*-i2 and *nad4*-i2 in *nmat1* (Keren, *et al*. 2012). The steady-state levels of several other intron-containing pre-mRNAs, in particularly *cox2*, *nad1*, *nad5* and *nad7*, were also higher in *nmat2* mutants than in wild-type plants (Keren et al., 2009; Zmudjak et al., 2017). Yet, the m^6^A profiles of various intronless transcripts in *nmat1* or *nmat2* mitochondria (Fig. 9), did not any show significant differences compared with the wild-type (col-0) plants. These results suggest that organellar transcripts affected in their metabolism may be more prone to *N^6^*-adenosine methylations.

**Figure 9.**
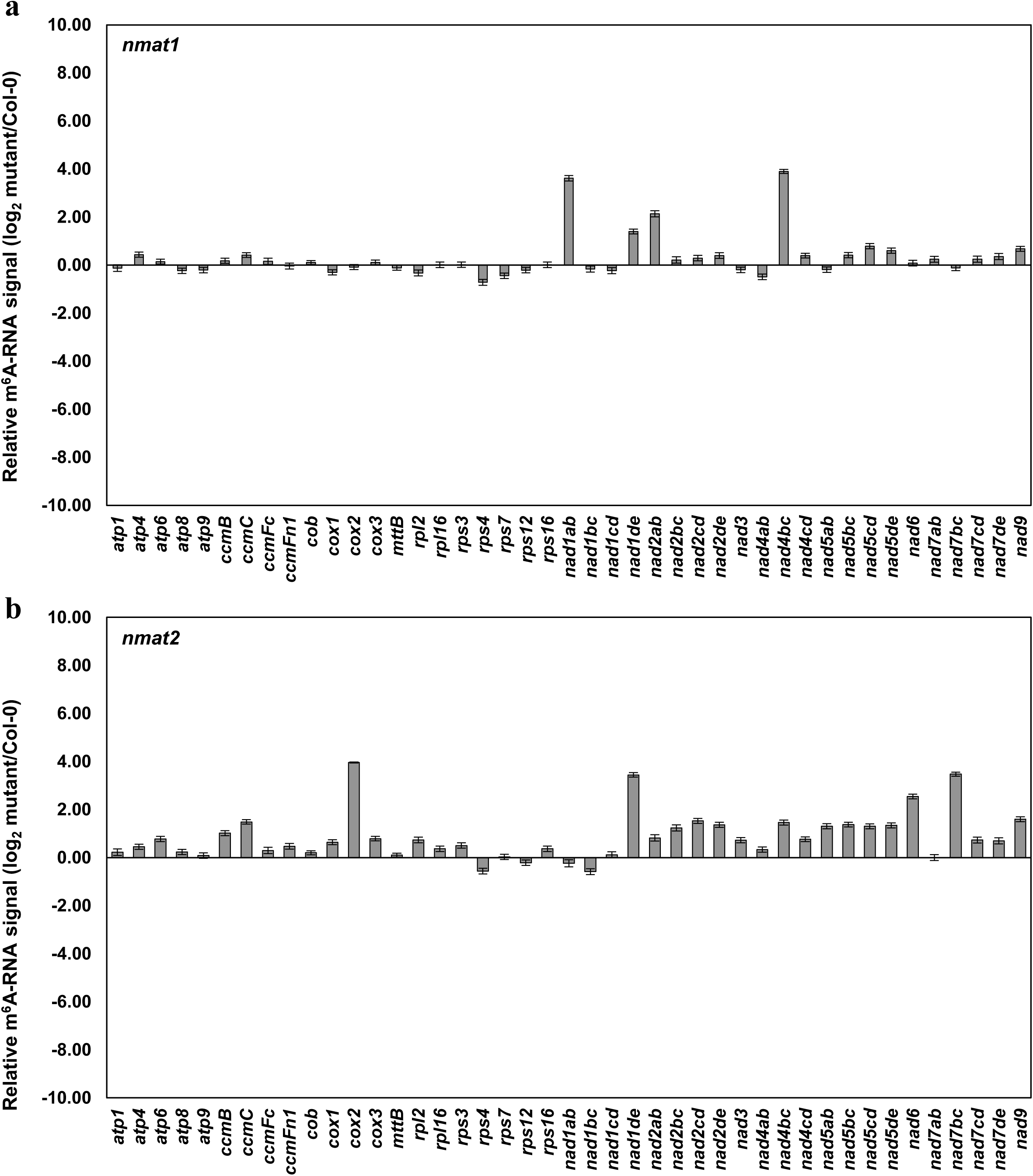
m^6^A methylomes of Arabidopsis mutants affected in mtRNA metabolism. The m6A methylation landscapes of *nmat1* (a) and *nmat2* (b) were analyzed by RT-qPCR following m6A pulldown assays, using anti-m6A antibodies. Total RNA extracted from wild-type (Col-0) and *nmat1* (Keren, *et al*. 2012, Nakagawa and Sakurai 2006) and *nmat2* (Keren, *et al*. 2009, Zmudjak, *et al*. 2017) mutants was used in m^6^A-RIP assays, with monoclonal m^6^A antibodies. The relative accumulation of different m^6^A-modified RNAs was assayed by RT-qPCR analyses with oligonucleotides corresponding to different genes in *Arabidopsis thaliana* mitochondria (Table S5). Panels ‘a’ and ‘b’ represent the relative m^6^A-RNA abundances in *nmat1* and *nmat2* mutants versus wild-type plants, respectively. Data in panels ‘a’ and ‘b’ represent the mean (± standard deviation, SD) of 15 replicates in 5 independent experiments (i.e., each performed in triplicate, and are presented relative to control).

**Figure 10.**
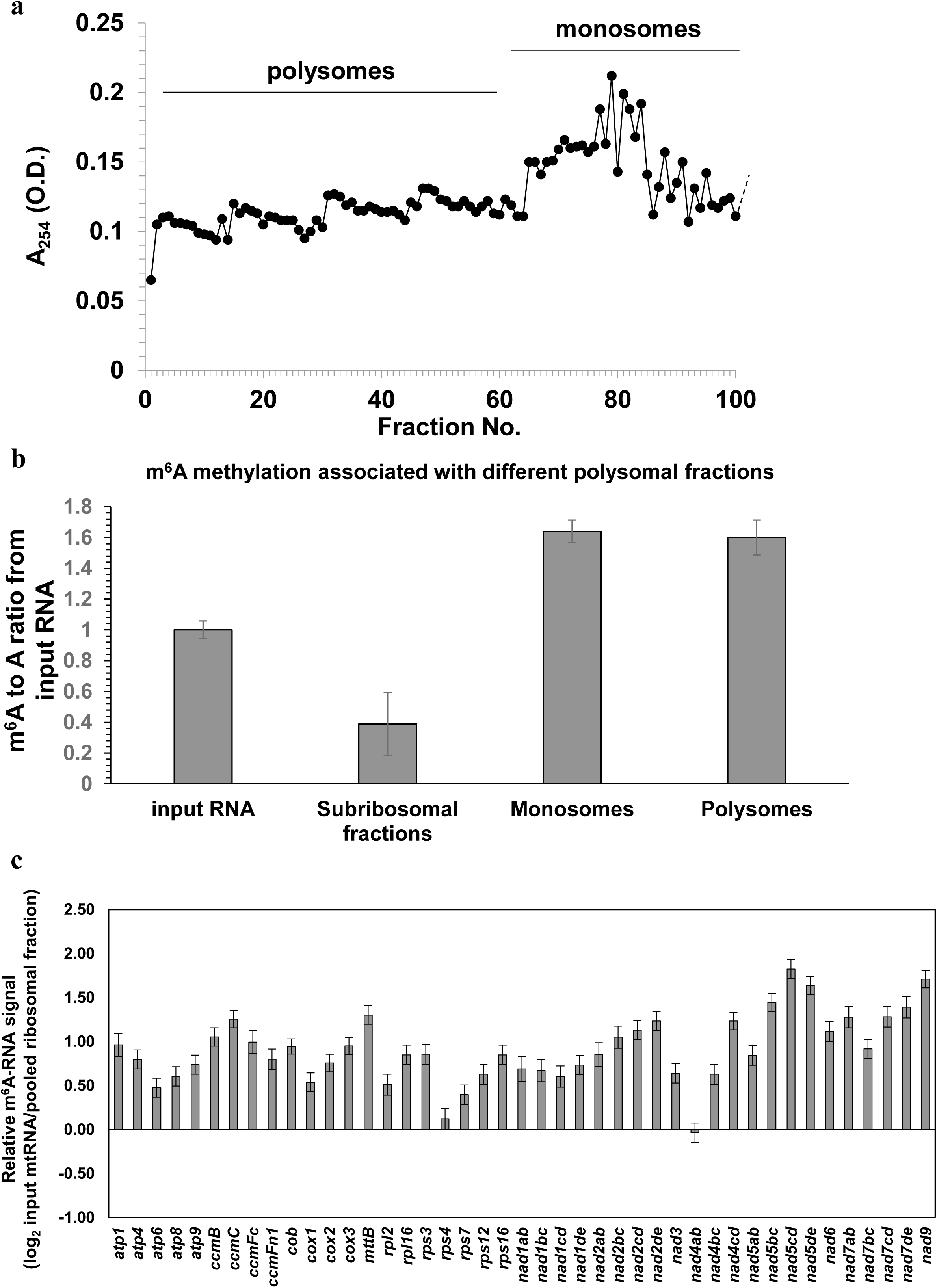
m^6^A-mRNA methylation associated with different polysomal fractions. Mitochondria isolated from cauliflower inflorescences were fractionated by sucrose gradient centrifugation. (a) Polysome profiles of cauliflower mitochondria (collected from bottom to top). (b) m^6^A mRNA extracted from different fractions quantified by calorimetric assay. (c) The relative levels of m^6^A-RNAs versus the input mRNA, evaluated by RT-qPCR of various mitochondrial mRNAs following m^6^A pulldown assays, using anti-m^6^A antibodies (i.e., log2 ratios). Data represent the mean (± standard deviation, SD) of 12 replicates in 4 independent experiments (i.e., each performed in triplicate, and are presented relative to control). Input correspond to ribosomal-depleted mtRNA before fractionation procedures, while fractions 105-150 indicate the sub-ribosomal fractions, fractions 65-95 correspond to mitochondrial monosomes, and fractions 5-50 relate to the organellar polysomes. Error bar shows standard deviation (STDEV) values. Each measurement was the average of three biological replicates. Small italic letters in panel ‘c’ indicate a significant difference from input RNA values (Student’s T-test, P 0.05).

### m^6^A mRNAs are enriched to some extent in the mitoribosomes

Published data (Akichika, *et al*. 2019, Choi, *et al*. 2016, Hoernes, *et al*. 2016, Meyer and Jaffrey 2017, Meyer, *et al*. 2015, Slobodin, *et al*. 2017, Sun, *et al*. 2019) and own results (Figs. 7 and 8a) indicate that the posttranscriptional modification of m^6^A can affect the translation of both nuclear and organellar-encoded polypeptides. The reduction in translation efficiency in the WGE system is correlated with magnitude of *N^6^*-adenosine modifications within a given transcript, while a single (or a few) m^6^A modification near initiation codons may enhance in some cases the translatability of the organellar transcripts. Polysomal-profiling experiments of purified cauliflower mitochondria were used to identify m^6^A-mRNA that are stably bound to the mitoribosomes (Waltz *et al*. 2019) (see Materials and Methods).

Figure 9a shows a typical polysome profile of cauliflower mitochondria. The fractions were pooled (bottom to top) into the heavier polysomal fractions (tubes 5 – 50) and the lighter monosomal fractions (tubes 65 – 95). In addition to the polysomal fractions, we also analyzed the m^6^A profiles in small-RNPs or subribosomal fractions (tubes 105 – 150), in comparison with those of the input RNA (rRNA-depleted). As shown in Figure 9b and Table S2, an enrichment in the m^6^A signals was apparent in both the ‘monosomal’ and ‘polysomal’ fractions. The relative levels of various mitochondrial transcripts in the polysomes (i.e., monosomal and polysomal fractions) were assayed by m^6^A-RIP followed by RT-qPCR analyses, using the input RNA (ΔrRNAs) as calibrator (Fig. 9c). The data indicated to small, yet statistically significant, increases (between 2x and 4x) in the levels of ribosome-associated m^6^A-mRNAs versus their ratios in the input RNA (Fig. 9c).

## Discussion

The *N^6^*-methyladenosine (m^6^A) was reported in the early 1970s (Desrosiers *et al*. 1974), and is currently considered as the most prevalent modification in nuclear-encoded mRNAs. co-IPs coupled to high-throughput RNA sequencing methods (Dominissini, *et al*. 2012), provided with comprehensive data regarding the m^6^A landscapes across the transcriptomes of various organisms (Burgess, *et al*. 2016, Liu and Pan 2016, Parker, *et al*. 2019, Wang, *et al*. 2017, Yue *et al*. 2015). Unlike pseudouridylation (U→Ψ), the most abundant posttranscriptional modification in RNAs (Ge and Yu 2013), or C→U exchanges, which are highly common in plant organelles (Barkan and Small 2014, Chateigner-Boutin and Small 2010, Colas des Francs-Small and Small 2014, Hammani and Giege 2014, Ichinose and Sugita 2016a, Schallenberg-Rüdinger and Knoop 2016, Shikanai 2015, Zmudjak and Ostersetzer-Biran 2017), m^6^A is not expected to rewire the genetic-code for translation (Dai *et al*. 2007). Instead, the m^6^A-modification is anticipated to serve important regulatory roles in gene expression in eukaryotic and prokaryotic organisms (see *e.g.*, (Bodi, *et al*. 2012, Deng, *et al*. 2015, Dominissini, *et al*. 2012, Hoernes, *et al*. 2016).

In bacteria, many of the m^6^A sites are mapped to genes of the respiratory apparatus (Deng, *et al*. 2015). These data may suggest an important regulatory role for m^6^A in controlling the OXPHOS capacity (Deng, *et al*. 2015), an activity which may have retained in the energy producing organelles (i.e., mitochondria and chloroplasts) in eukaryotic systems. Global transcriptomic analyses indicated the presence of m^6^A-modifications in mitochondrial (and plastidial) transcripts in the model plant, *Arabidopsis thaliana* (Bodi, *et al*. 2012, Chen and Liu 2014, Luo, *et al*. 2014, Shen, *et al*. 2016, Wang, *et al*. 2017). However, as the RNA-seq analyses involved total plant RNA preparations, the occurrence and significance of *N^6^*-methyladenosine modifications within the organelles in plants (and animals) need to be further investigated. In this study, we provide with detailed analyses of the m^6^A-RNA landscapes in the mitochondria of two *Brassicaceae* species, i.e., Arabidopsis and cauliflower.

Northern blots (Fig. S1), calorimetric assays (Table S2) and LC-MS analyses (Figs. 2 and S2), indicated the presence of m^6^A within organellar transcripts in Arabidopsis and cauliflower mitochondria, with a calculated m^6^A occurrence of about 0.4 to 0.5% of the total organellar A residues, on average. Following the confirmation of m^6^A modification in plants organellar transcripts, the topologies of m^6^A-RNA methylomes in the mitochondria of Arabidopsis and cauliflower were analyzed by the m^6^A-RIP-seq method (Fig. 1, and (Dominissini, *et al*. 2012, Luo, *et al*. 2014).

The m^6^A-RIP-seq analyses indicated to different m^6^A modification patterns in Arabidopsis and cauliflower mitochondria (Fig. 3, Tables 1, S3 and S4). Such differences could relate to the analyses of different plant species, altered growth conditions, or may indicate differences in m^6^A-RNA modifications patterns between different tissues (i.e., whole plant versus inflorescences). Despite the differences, the m^6^A-RIP-seq of Arabidopsis and cauliflower mitochondria generally showed a similar distribution of the m^6^A sites, where about half of the reads are mapped to noncoding RNAs and about quarter (i.e., 23% in Arabidopsis seedlings and 27% in cauliflower inflorescences) of the m^6^A signals are found in known protein-coding genes (Fig. 3a and Table 1). While in nuclear-encoded transcripts in animals and plants, the majority of the m^6^A sites are mapped to regions adjacent to stop codons within the 3’ UTRs (Bodi, *et al*. 2012, Deng, *et al*. 2015, Dominissini, *et al*. 2012, Hoernes, *et al*. 2016), in Arabidopsis and cauliflower mitochondria, the majority of the m^6^A sites within known genes are mapped to the coding regions (i.e., 60.8% in Arabidopsis and 52.3% in cauliflower). In fact, less than 5% of the m^6^A reads are mapped to UTRs in both Arabidopsis and cauliflower mitochondria (Fig. 3a and Table 1). These data also correlate with the distribution of m^6^A in the mtRNAs of Arabidopsis by the global RNA-seq analyses s (Luo, *et al*. 2014). Notably, a significant portion of m^6^A reads were found to reside within group II intron sequences (i.e., 17% in Arabidopsis and 18% in cauliflower) (Figs. 3, S3 and Table 1).

Beside to their unique topology, plant mitochondria also exhibit a distinctive distribution of m^6^A modifications within their transcripts. While noncoding RNAs contain multiple m^6^A sites distributed along the transcript (Figs. 3b and S4), the coding sequences contain a fewer number of m^6^A-modifications per transcript, with a notable preference to regions close (±100 nts) to the AUG sites (i.e., 27.3% in Arabidopsis and 12.0% in cauliflower) (see Fig. 3b). These data may indicate that the m^6^A-modification plays a regulatory role in the expression of organellar genes in angiosperms mitochondria. The m^6^A-RIP-seqs data also revealed the presence of various RNA reads which mapped to the plastid genome (see supplementary Table S6 and Fig. S6). However, these data represent preliminary data and the topology of m^6^A modifications in the plastids need to be further investigated.

Through its impact on splicing (Xiao *et al*. 2016), mRNA stability (Wang *et al*. 2014) and rate of translation in animals (Meyer, *et al*. 2015, Wang *et al*. 2015b), m^6^A regulate essential features of a cell by affecting the expression of nuclear-encoded genes. In Arabidopsis mutants, altered m^6^A methylation patterns result with embryogenesis and developmental defect phenotypes (Bodi, *et al*. 2012, Luo, *et al*. 2014, Růžička *et al*. 2017, Shen, *et al*. 2016, Zhong, *et al*. 2008). It is therefore anticipated that m^6^A would also affect the expression of organellar genes.

Following the analysis of the topologies of m^6^A in Brassicales mitochondria, we next sought to infer the roles of m^6^A-modifications in organellar biogenesis and gene-expression. As we were unable to determine the significance of m^6^A to mitochondria biology by forward or reverse genetis, we used an in vitro approach to study the effects of *N^6^*-adenosine methylations on organellar gene expression, using the wheat germ extract (WGE) system. No significant effects on the stability of recombinant *atp1*, *nad4* or *matR* transcripts was observed between unmodified and m^6^A-modified transcripts in vitro (Figs 6 and S5). However, excessive m^6^A modifications were found to be detrimental for the translation of the m^6^A-modified transcripts in vitro (Fig. 7). This effect are related to organellar translation, as the translatabilty of ATP1, NAD4 and MATR were reduced in the presence of cycloheximide, which affects the translation of mRNAs bound to cytosolic ribosomes (Fig. 7a,b).

We further analyzed the effect of a single modification within the start (AUG) codon on the translation efficiencies of ATP1, MATR and NAD4 fragments, using synthetically modified RNAs. While reduced translation was observed in the case of ‘body-labeled’ *atp1*, *nad4* and *matR* RNA (i.e., modified at multiple sites along their transcripts) (Figs. 5 and 7), the translation efficiencies of m^6^AUG-modified transcripts increase, at least to some extent, in the case of *atp1* and *nad4* (Fig. 8a). Also, no significant effects on the translatability of *atp1*, *matR* and *nad4*, modified at the initiation codon, was seen in the presence of Kanamycin, which affects the translation of organellar proteins (Wang, *et al*. 2015a). Together, these results suggest that multiple m^6^A modifications are detrimental for translation, while a single modification in the start codon (and maybe also of other A residues found in proximity to the translation imitation site), can enhance the translation of some organellar transcripts. The molecular basis for these effects are currently unknown. It was previously shown that posttranscriptional m^6^A-modifications affect the translation of various nuclear-encoded transcripts (Akichika, *et al*. 2019, Choi, *et al*. 2016, Hoernes, *et al*. 2016, Meyer and Jaffrey 2017, Meyer, *et al*. 2015, Slobodin, *et al*. 2017, Sun, *et al*. 2019). In a recent study, (Akichika, *et al*. 2019) indicated that the translation efficiency of some mRNAs in yeast, are significantly decreased upon KO of *CAPAM* gene a cap-specific adenosine-*N^6^*-methyltransferase. It is possible that an organellar writer protein for *N^6^*-methylation may possess similar functions. Following the modifications in sites adjacent to the translation start sites, specific readers may associate with the modified transcripts and enhance their translatability.

Under the *in vitro* conditions, the transcription efficiency of synthetic RNA-fragments corresponding to *atp1*, *nad4* and *matR* was not significantly affected by m^6^A, even at m^6^A/A ratio’s as high as 50% (Fig. 5). A reduction in the transcription efficiencies of cDNAs corresponding to *atp1*, *nad4* and *matR* was observed only when the reaction mix contains a 100% m^6^A/A ratio (Fig. 5). It is expected that m^6^A modifications occur posttranscriptionally. Accordingly, it was recently shown that both plant and animal cells require an *N^6^*-mAMP deaminase (MAPDA) to catabolize *N^6^*-mAMP to inosine monophosphate, most likely to prevent misincorporation of m^6^A into the RNAs during transcription (Chen *et al*. 2018). This is further supported by *in vitro* assays that show that eukaryotic RNA polymerases (RNAPs) can also use *N^6^*-mATP as a substrate (Chen, *et al*. 2018). As, the single-subunit RNA polymerase of plant mitochondria (mtRNAP) is related to the T7/T3 RNAPs (Cermakian *et al*. 1996, Masters *et al*. 1987, Zmudjak and Ostersetzer-Biran 2017), we speculate that the incorporation of m^6^A may precedes efficiently in the presence of m^6^ATP ribonucleotides in the mitochondria *in vivo*, as well (unless *N^6^*-mAMP is catabolized in these oraganlles).

Examining common m^6^A motifs in Arabidopsis and cauliflower mitochondria, using the HOMER (Heinz, *et al*. 2010) and MEME (Bailey, *et al*. 2009) programs, we observe a statistical enrichment for the 5’-Rm^6^AY-3’ sequence (P<0.005; Fig. 4). The predicted mitochondrial m^6^A sites shares homology with the m^6^A core motif suggested for nuclear-encoded mRNAs (i.e., RR<u>m^6^A</u>CH) in animals and plants (Liu and Pan 2016, Maity and Das 2016, Patil, *et al*. 2018), and to a lesser degree with the m^6^A motif of bacterial RNAs (*i.e*., UGC<u>Cm^6^AG</u>). However, the identities of the enzymes which regulate m^6^A patterns in plant mitochondria remain unknown.

The nuclear genome of *Arabidopsis thaliana* contains several genes which are closely related to m^6^A methyltransferases (*i.e*., Writers) in mammals (see *e.g*. (Růžička, *et al*. 2017) and Table S7). These include an MTA-related protein (At4g10760) and an ortholog of the mammalian WTAP (AT3G54170), known as FIP37. The two factors serve as core components of the m^6^A methyltransferase complex, and play indispensable roles in determining the m^6^A mRNA modification patterns in Arabidopsis plants (Bodi, *et al*. 2012, Luo, *et al*. 2014, Růžička, *et al*. 2017, Shen, *et al*. 2016, Zhong, *et al*. 2008). Here, we performed sequence similarity searches with PSI-BLAST, in an effort to identify putative components of the organellar m^6^A methyltransferase complex from Arabidopsis and rice genomes. These analyses revealed the existence of several genes which share homology with known m^6^A enzymes, are predicted to reside with the mitochondria or chloroplasts, and are conserved between dicot and monocot species (Table S7). Among the factors which match such criteria are At-MTA and FIP37 proteins, which were also identified in proteomic analyses of plant organellar preparations (see Table S7). These observations may also relate to the similarities in the m^6^A sequence motifs found between the nuclear- and organellar-encoded transcripts (Fig. 4b). The location and functions of candidate factors (*e.g.*, At-MTA and FIP37) in m^6^A-RNA modifications in plant organelles are currently under investigation.

### Conclusions

The expression of the mitochondrial genomes in plants is complex, particularly at the posttranscriptional level. RNA processing events that contribute to mitochondrial gene expression include the processing of polycistronic transcripts, trimming, splicing and deamination of many Cytidine residues to Uracil (reviewed in e.g., (Zmudjak and Ostersetzer-Biran 2017). More than 100 types of RNA-base modifications are currently known (Nachtergaele and He 2017). These RNA modifications can affect cellular activities by altering the genetic code (*e.g*., pseudouridylation and deamination), affecting the RNA structure, or by recruiting specific cofactors (readers) that influence a wide range of biological functions, e.g., transcription, stability, processing and translatability. The m^6^A modifications show dynamic patterns, changing in their distribution and levels in response to various environmental or developmental signals. Here, we show that plant mitochondrial RNAs undergo many *N^6^*-adenosine methylation (m^6^A) modifications in different RNA types (*i.e*., intergenic-expressed transcripts, tRNAs, mRNAs, intron and UTR sequences), with an occurrence of about 5 m^6^A sites per 1,000 Adenosine residues. Does m^6^A affects organellar gene expression in plants? Currently, we cannot provide a definitive answer. The in vitro translation assays may indicate that m^6^A have a selective program: it is detrimental for the translation of RNAs modified at multiple sites (e.g., as introns and noncoding RNAs), while methylations within, or near, the start codon may rather enhance the translation. Based on the data, we speculate that m^6^A may adds an additional layer of posttranscriptional regulation in the expression program of plant mitochondria, which play critical roles in cellular metabolism. Accordingly, m^6^A seems enriched in ribosome-associated mRNAs, while mutants affected in organellar gene expression show specific changes in their m^6^A methylomes. However, such assumptions need to be supported experimentally. Analysis of Arabidopsis plants affected in genes that function in m^6^A modification in plant mitochondria would provide with more insights into the roles of m^6^A modifications to mitochondria biogenesis and gene expression. We further intend to define whether the m^6^A landscapes in plant mitochondria change in response to physiological and environmental stimuli.

## Experimental procedures

### Plant material and growth conditions

Plants growth and analyses generally followed the procedures described in (Sultan, *et al*. 2016). *Arabidopsis thaliana* ecotype Columbia (Col-0) seeds were obtained from the ABRC center, at Ohio State University (Columbus, OH). The seeds were sown on MS-agar plates, incubated in the dark for 2 days at 4°C, and then transferred to controlled temperature and light conditions in the growth chambers [*i.e*. 22°C under short day conditions (8 hrs. light, 150 µE·m-2·s-1 and 16 hour dark)]. Cauliflower (*Brassica oleracea* var.) inflorescences were purchased fresh at local markets and used immediately.

### Mitochondria extraction and analysis

Isolation of mitochondria from 3-week-old Arabidopsis seedlings grown on MS-plates was performed according to the method described in (Keren, *et al*. 2012). Preparation of mitochondria from cauliflower inflorescence followed the protocol described in detail in (Neuwirt, *et al*. 2005). In brief, about 1.0 kg cauliflower inflorescences were cut off from the stems, and ground with ice-cold extraction buffer [0.9 M mannitol, 90 mM Na-pyrophosphate, 6 mM EDTA, 2.4% PVP25 (w/v), 0.9% BSA (w/v), 9 mM cysteine, 15 mM glycine, and 6 mM β-mercaptoethanol; pH 7.5]. Mitochondria were recovered from the extract by differential centrifugations and purification on Percoll gradients, aliquoted and stored frozen at −80°C.

### RNA extraction and analysis

RNA extraction from highly-purified mitochondria was performed essentially as described previously (Cohen *et al*. 2014, Keren, *et al*. 2011, Zmudjak, *et al*. 2017). In brief, RNA was prepared following standard TRI Reagent^®^ protocols (Molecular Research Center Inc., TR-118). The RNA was treated with RNase-free DNase (Ambion, Cat no. AM2222) prior to its use in the assays. Reverse transcription was carried out with Superscript III reverse transcriptase (Thermo Fisher Scientific, Cat no. 18080093), using 1 μg of total Arabidopsis or cauliflower mtRNA and 250 ng of a random hexanucleotide mixture (Promega, Cat no. C1181) and incubated for 50 min at 50°C. Reactions were stopped by 15 min incubation at 70°C. Quantitative reverse transcription PCR (RT-qPCR) was performed essentially as described previously (Sultan, *et al*. 2016, Zmudjak, *et al*. 2017).

### Detection of *N^6^*-Adenosine methylations (m^6^A) in mitochondrial RNAs by LC-UV-MS/MS

Detection of m^6^A methylations in mitochondrial RNA (mtRNA) was carried out by LC-UV-MS/MS, essentially as described previously (Thuring *et al*. 2016). In brief, 75 μg total mtRNA was digested to single nucleosides, using 10 U S1 nuclease (Thermo Fisher Scientific, Cat no. EN0321), 1 U snake venom phosphodiesterase (Exo-PI; Sigma-Aldrich, Cat no. P4506), and 10 U alkaline phosphatase (Thermo Fisher Scientific, EF0654). The nucleosides products were then analyzed by liquid chromatography-mass spectrometry, using Luna^®^ polar (RP-C18) column (Phenomenex, Aschaffenburg, Germany). Data were acquired and processes using the MassHunter software (version B.06.00, Agilent Technologies, Wilmington, DE, USA). %Table S1 summarizes the chromatographic data, including the retention times (RT), masses, abundances and molecular formula of the analyzed products.

### Calorimetric m^6^A measurements

Global m^6^A modifications in total organellar RNAs was assessed by the EpiQuik m^6^A RNA Methylation Quantification Kit (Epigentek Group Inc., Farmingdale, NY) according to the manufacturers’ specifications, with 200 ng total RNA, in duplicates.

### m^6^A-RNA co-immunoprecipitation (m^6^A-RIP)

Monoclonal anti-m^6^A antibody (Synaptic systems, Cat no. 202011) was incubated with Protein A/G Magnetic Beads (Pierce, 88802) for 30 min at room temperature in IPP buffer (0.2 M NaCl, 0.1% NP-40, 10 mM Tris-HCl, pH 7.5, 1 U/μl RNasin). The antibody-bound beads were washed twice with IPP buffer and resuspended in 200 µl of IPP buffer. Prior to m^6^A-RIP, the ribosomal RNA was removed from the total mtRNA using the RiboMinus™ Plant Kit for RNA-seq (Thermo Fisher Scientific, Cat no. A1083808). The purified organellar RNA was then fragmented using an RNA Fragmentation kit (Ambion, Cat no. AM8740), yielding fragments in the range of 60 to 200 nts. The fragmented RNA was denatured at 70°C for 2 min and then incubated for 2∼4 hrs. at 4°C with 5 μg anti-m^6^A antibody coupled to magnetic protein A/G beads, in the presence of RNase inhibitor (Ambion, Cat no. AM2682). The beads were then washed twice with IPP buffer, twice with low salt buffer (50 mM NaCl, 0.1% NP-40, 10 mM Tris-HCl pH 7.5), twice with high salt buffer (0.5 M NaCl, 0.1% NP-40, 10 mM Tris-HCl pH 7.5) and once with IPP buffer. The m^6^A-RNA was released from the beads with RLT buffer (Qiagen, 79216) and purified using the Dynabeads^TM^ MyOne^TM^ Silane RNA isolation Kit (Thermo Fisher Scieintific, Cat no. 37002D). For the analysis of m^6^A in mtRNAs (i.e., cauliflower) we further used a non-radioactive northern blotting (Keren, *et al*. 2011), with anti-m^6^A antibodies (Fig. S1).

### Transcriptome mapping by high-throughput (RNA-seq) analysis

Total mtRNA was extracted from mitochondria preparations, obtained from Arabidopsis and cauliflower inflorescence. Sequencing was carried out on an Illumina Genome Analyzer (The Center for Genomic Technologies, The Hebrew University of Jerusalem, Israel), generally as described previously (Grewe, *et al*. 2014). Reads were first filtered for low quality and primers or adapters contaminations using trimomatic (version 0.36) (Bolger *et al*. 2014). Filtered reads were mapped to the *A. thaliana* and *B. oleracea* mitochondrial genomes (NC_001284 and KJ820683.1, respectively) using Bowtie2 (version 2.2.9) (Langmead and Salzberg 2012). BEDTools (Quinlan and Hall 2010) was used to determine the coverage of mapped reads of the m6A-RIP-seq and input RNA RNA-seq libraries on the genomes in a 1 bp intervals, and m6A peaks were called using an R-script modified from (Luo, *et al*. 2014) (Supplementary script 1). Only significant peaks (p-value < 0.05) were selected for further analysis (Supplementary datasets 1&2). Resulting m6A peaks were mapped to annotated genes using BEDTools intersect. Consensus of methylation sites was predicted using HOMER (Heinz, *et al*. 2010). Sequences are available at the Sequence Read Archive (SRA, accession PRJNA472433).

### Synthesis of recombinant His-tagged mitochondrial proteins

*In vitro* transcription assays were performed as described previously (Ostersetzer and Adam 1997, Ostersetzer and Adam 1999). Gene-fragments corresponding to ATP1, MatR and NAD4 were generated by RT-PCRs with specific oligonucleotides designed to each gene-fragment (i.e., containing the *Nco* I and *Xho* I restriction sites) (supplementary Table S8). The fragments were digested with the restriction endonucleases *Nco* I and *Xho* I and cloned into same sites in pET28b (see Table S8). The integrity of each construct was confirmed by DNA-sequencing (Center for Genomic Technologies, HUJI, Israel). For *in vitro* transcriptions, 1 U of T7 RNA polymerase (Promega, Cat no. P2075) was added to 500 ng of the cDNA-fragments in T7 transcription reaction buffer (Promega, Cat no. P2075), and incubated for 2 hours at 37°C in the absence or presence of *N^6^*-methyladenosine (Jena Biosciences, Cat no. NU-1101L) to generate ‘body-labeled’ m^6^A-RNAs. mRNAs labeled in the start codon (*i.e*. m^6^AUG) were generated by custom synthesis (i.e., Tamar Laboratories, Israel, Integrated DNA Technologies, Inc., IDT), with the following sequence: TTTAA<u>GAAGGAGA</u>TATACC/**iN6Me-rA**/TG[xxx], where the ribosome binding (rbs, or Shine-Dalgarno) sequence is underlined, iN6Me-rA corresponds to the m^6^A modification (according to IDT), and the ‘xxx’ region corresponds to specific ATP1, MatR or NAD4 gene-fragments containing a 6x His-tag ligated to the synthetic m^6^AUG-RNA fragments (see Table S8). Recombinant His-tagged ATP1, MatR and NAD4 proteins were obtained by in vitro translation assays, using a coupled transcription-translation system containing wheat germ extract (TNT kit; Promega, Cat no. L5030), according to the manufacturer protocol.

### RNA stability assay

Transcript fragments corresponding to *matr*, *nad4* and *atp1* were transcribed *in vitro* with Biotin-14-CTP (ThermoFischer Scientific, 19519016) and in the absence or presence of m^6^ATP (see %Table S1). The *in vitro* transcribed RNAs were incubated with mitochondrial extracts for 2h at 30°C. For the analysis of RNA stability, organellar samples were taken at specific time points, as indicated in the blots, and RNA stability was analyzed by Filter binding assays (Barkan *et al*. 2007, Keren *et al*. 2008, Ostersetzer *et al*. 2005). Detection of RNAs was carried out by non-radioactive northern blots, using anti-Biotin antibodies and chemiluminescent detection.

### Isolation of ribosome-bound mRNAs from cauliflower mitochondria

Organellar polysomes were obtain generally as described previously (Barkan 1998, Kahlau and Bock 2008, Uyttewaal *et al*. 2008, Zoschke *et al*. 2013), with some modifications. Mitochondria isolated from cauliflower inflorescences were suspended in polysomal extraction buffer (*i.e.*, 40 mM Tris-acetate pH 8.0, 10 mM MgCl_2_, 0.2 M Sucrose, 10 mM β-mercaptoethanol, 1% Triton X-100, 2% polyoxyethylen-10-tridecylether, 0.5 mg/ml heparin, chloramphenicol 100 μg/ml, cycloheximide 25 μg/ml). The mitochondrial suspension was centrifuged (15,000 x g, 10 min at 4°C) to remove insoluble material, and the clear lysate was then loaded onto an step gradient containing 15%, 30%, 40% and 55% sucrose in polysome gradient buffer (0.4 M Tris-HCl pH 8.0, 0.2 M KCl and 0.1 M MgCl_2_) and centrifuged at 45,000 x g at 4 °C for 65 min at 4°C. Each gradient was then separated into several fractions (i.e., bottom-to-top) of equal volumes (about 100 μl) using a fraction collector (BioRad, Hercules, USA). RNA from all fractions was prepared following standard TRI Reagent® protocols (Molecular Research Center Inc., TR-118).

## Accession numbers

SRA accessions PRJNA472433

## Acknowledgements

We thank the Arabidopsis biological resource center for providing mutant seeds. The authors confirm that they have no conflict of interest to declare. This work was supported by grants to O.O.B from the ‘Israeli Science Foundation’ (ISF grant no. 741/15).

## Short legends for Supporting Information

**Supplementary script 1**. The R script used to in this study for m^6^A peaks discovery from RIP-RNA-seq data.

**Supplementary Data Set 1**. Arabidopsis RNA-seq data set.

**Supplementary Data Set 2**. Cauliflower RNA-seq data set.

---

**Figure S1.**
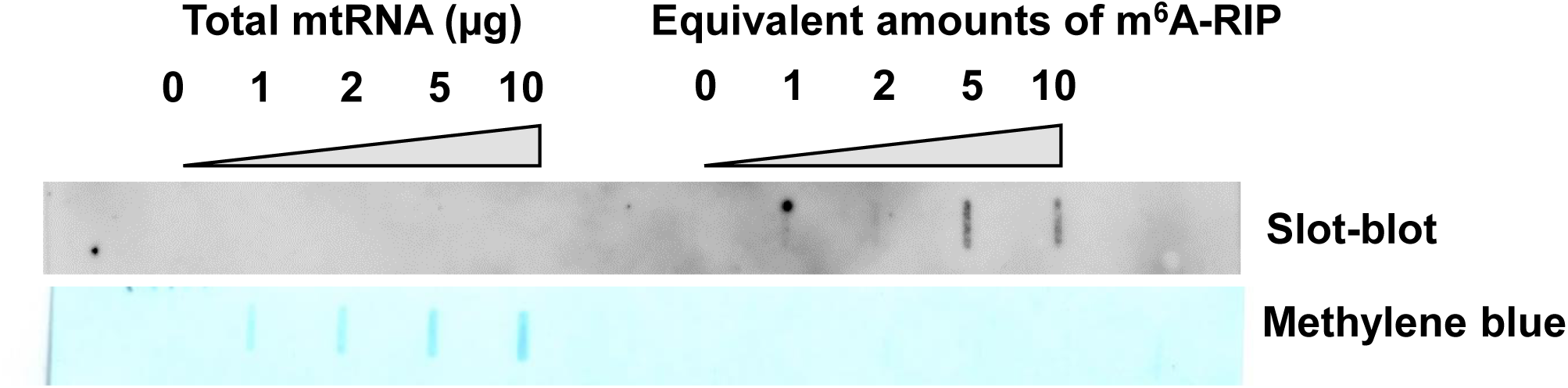
Analysis of m^6^A modifications in cauliflower mtRNA. The presence of m^6^A modifications was analyzed by slot blots and immunoassays with anti-m^6^A antibodies.

**Figure S2.**
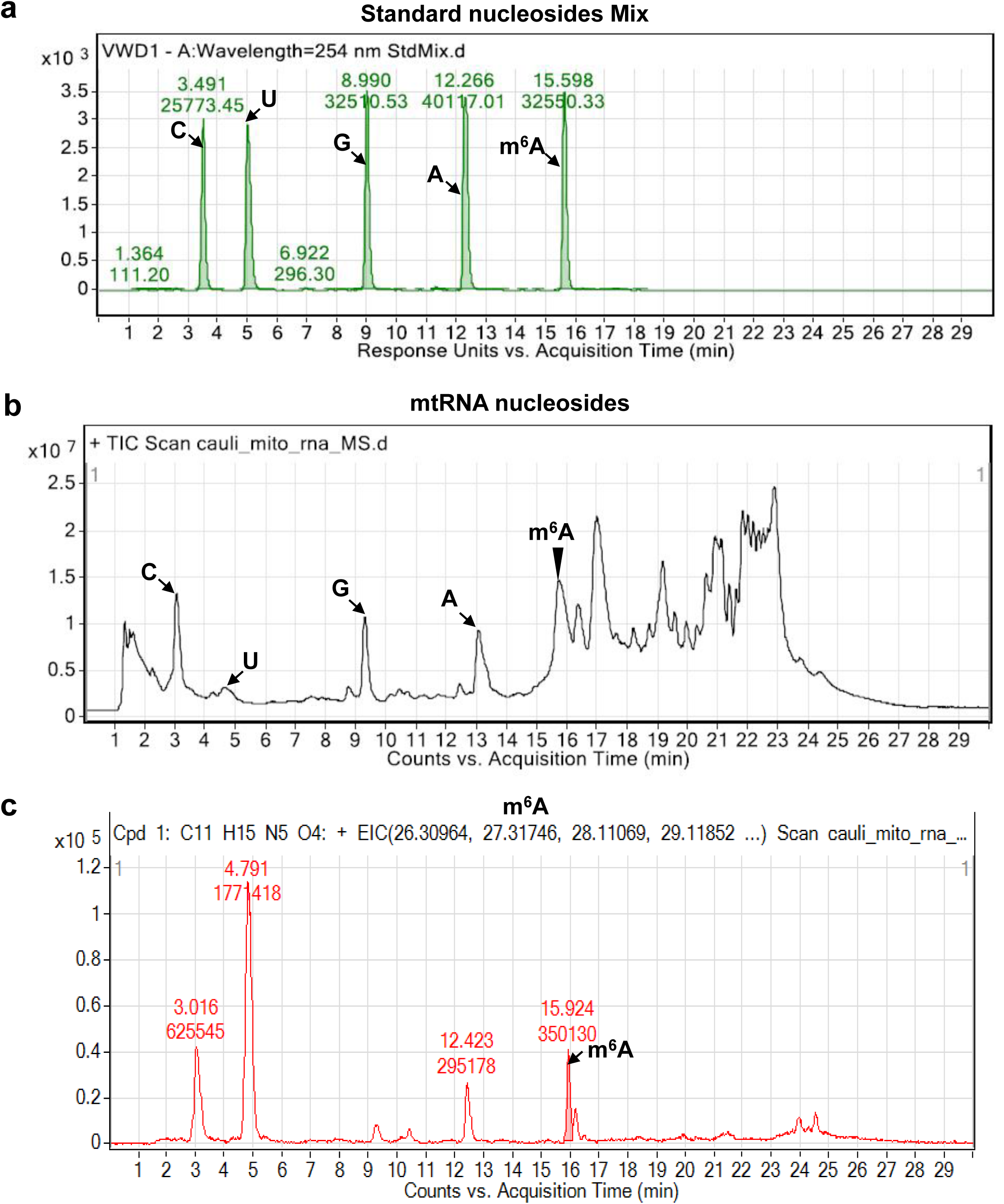
Identification of m^6^A by LC-MS/MS in mitochondrial RNAs isolated from wild-type cauliflower inflorescences. Total mtRNA, isolated from cauliflower mitochondria, was digested to single nucleosides with S1 nuclease and phosphodiesterase, and the presence of modified Adenosine residues (m6A) was assayed by LC-MS, based on specific retention times from standard and modified nucleosides separated by the same chromatographic procedure. The UV absorption spectra of (a) different nucleosides (A, G, C, and U) and m6A standards, (b) nucleosides obtained by in total RNA digestion and (c) the specific MS spectrum of m6A nucleosids in mitochondrial RNA preps are given in each panel.

**Figure S3.**
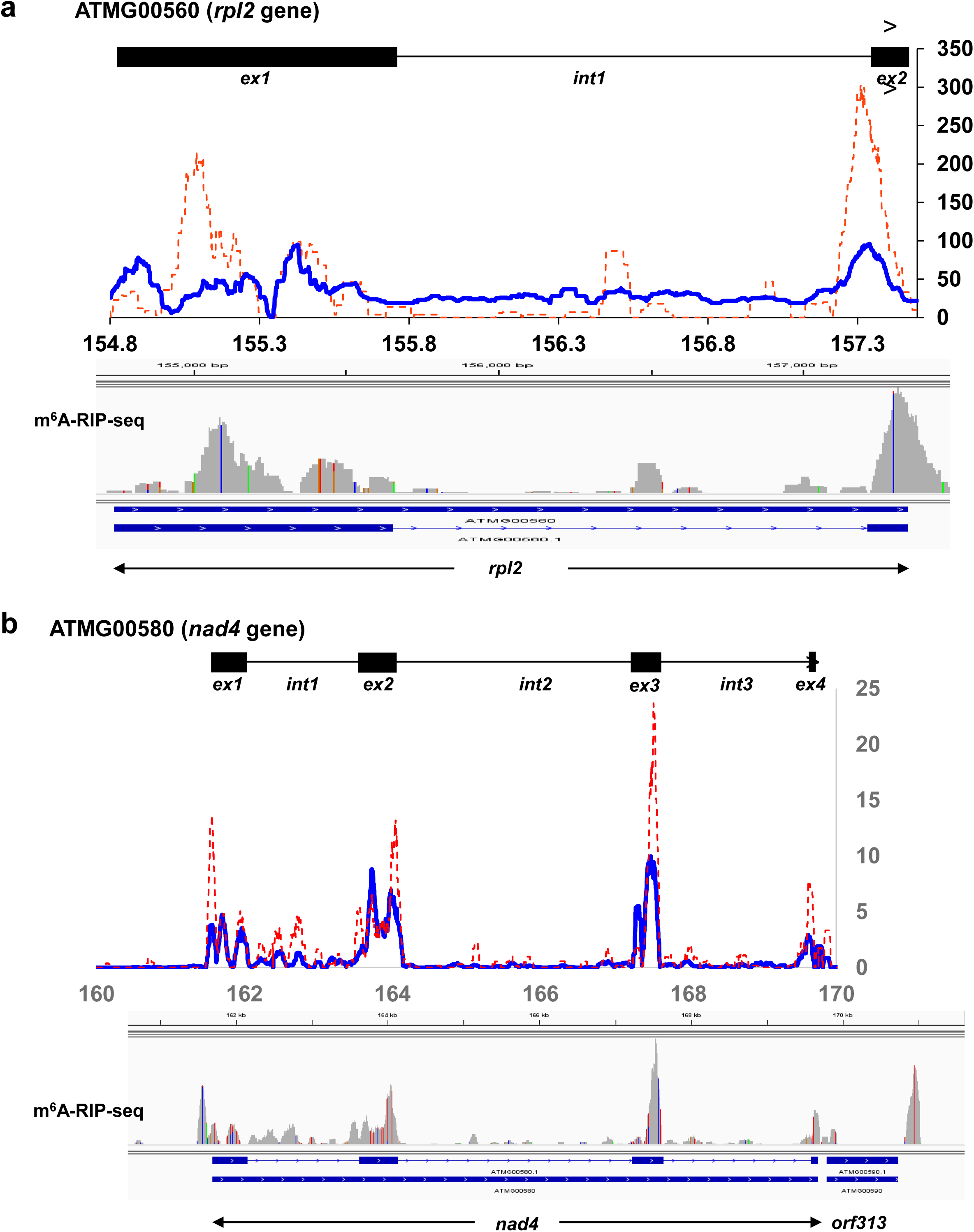
Representative analyses of *N^6^*-adenosine metylomes in Brassicales mtRNAs. The topology of m^6^A modifications was analyzed by m^6^A-RIP-seq analyses with anti-m^6^A antibodies. The reads were aligned to Arabidopsis mitochondrial genome (NC_001284.2; Unseld et al., 1997). Coverage of m^6^A (blue line) and control (input mtRNA, in red line) of two organellar genes, rpl2 (a) and nad4 (b) is indicated in the figure. To normalize for expression levels, coverage for each position is divided by the mean coverage in the control experiment.

**Figure S4.**
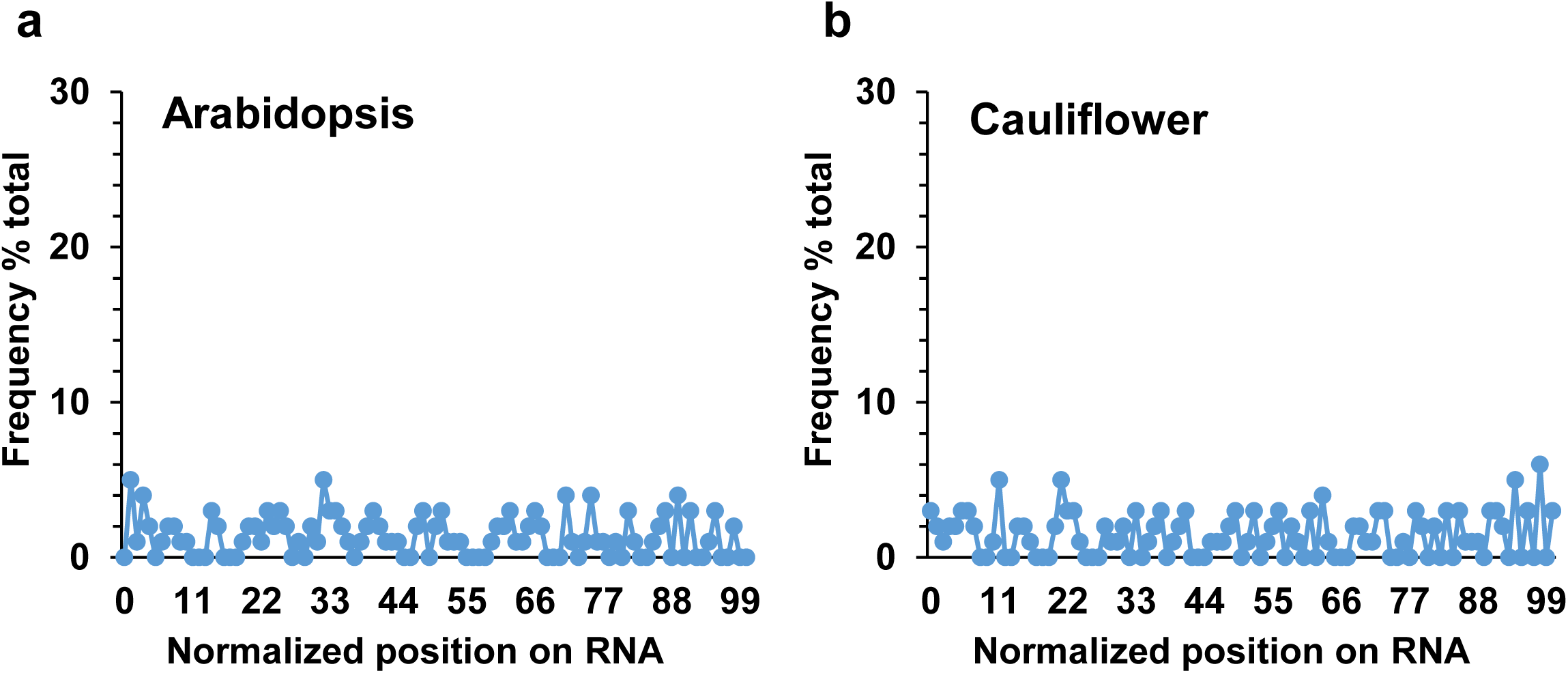
Distribution of m^6^A modifications in noncoding mtRNAs of Arabidopsis and cauliflower. Distribution of m6A peaks in Arabidopsis (a) or Cauliflower (b) sequences corresponding to noncoding RNAs, according to normalized gene lengths.

**Figure S5.**
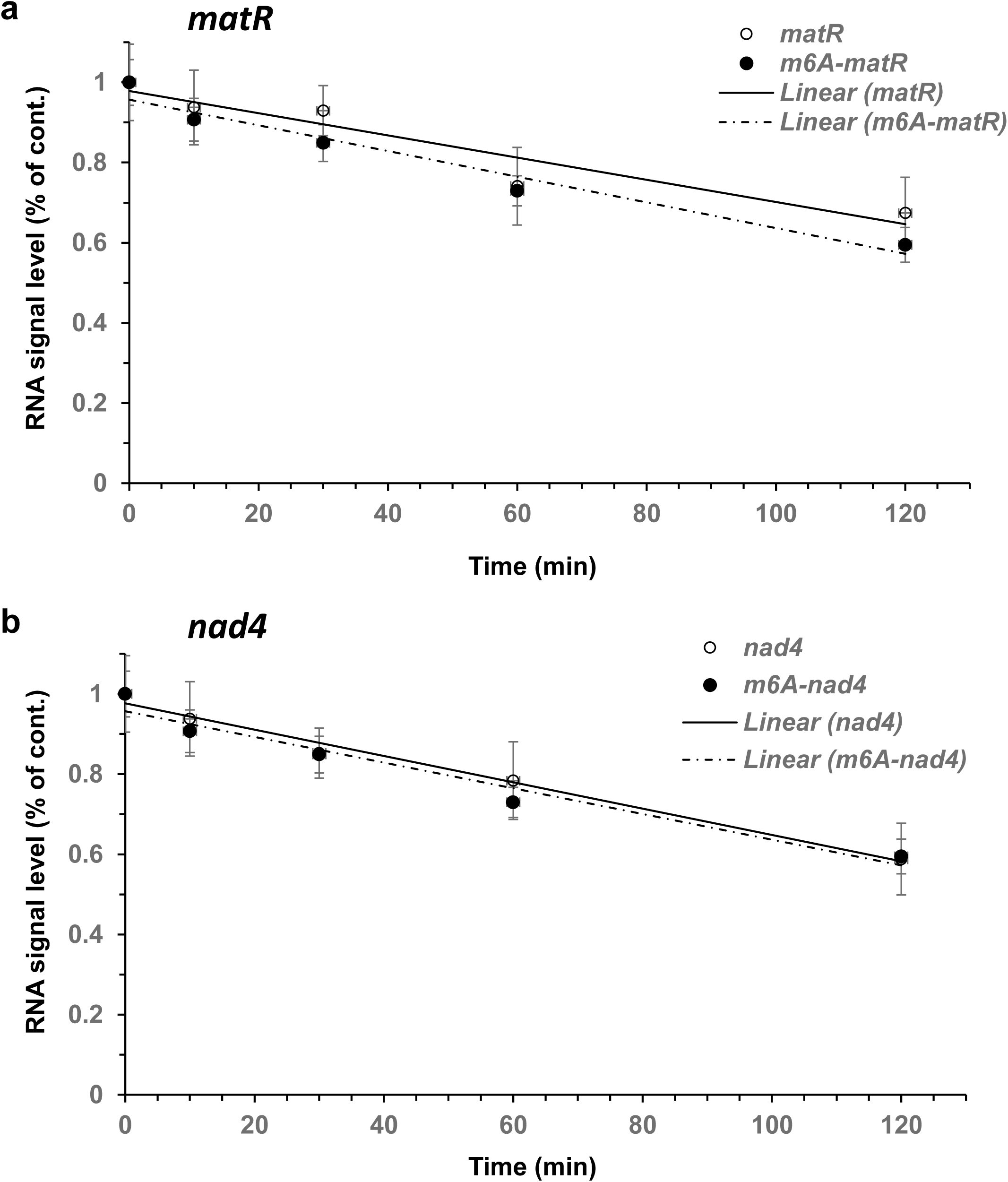
The effect of m6A modifications on mtRNA stability. The decay patterns of control (i.e., non-modified) versus m6A body labeled mRNAs corresponding to *matR* (a) or *nad4* (b), were examined by incubating the in vitro transcribed RNAs with purified cauliflower mitochondrial extracts. Samples were collected at different time points, as indicated in the figure, and the relative levels of *matR* and *nad4* mRNAs were examined by Filter binding assays (Ostersetzer et al., 2005; Keren et al., 2008; Keren et al., 2011).

**Figure S6.**
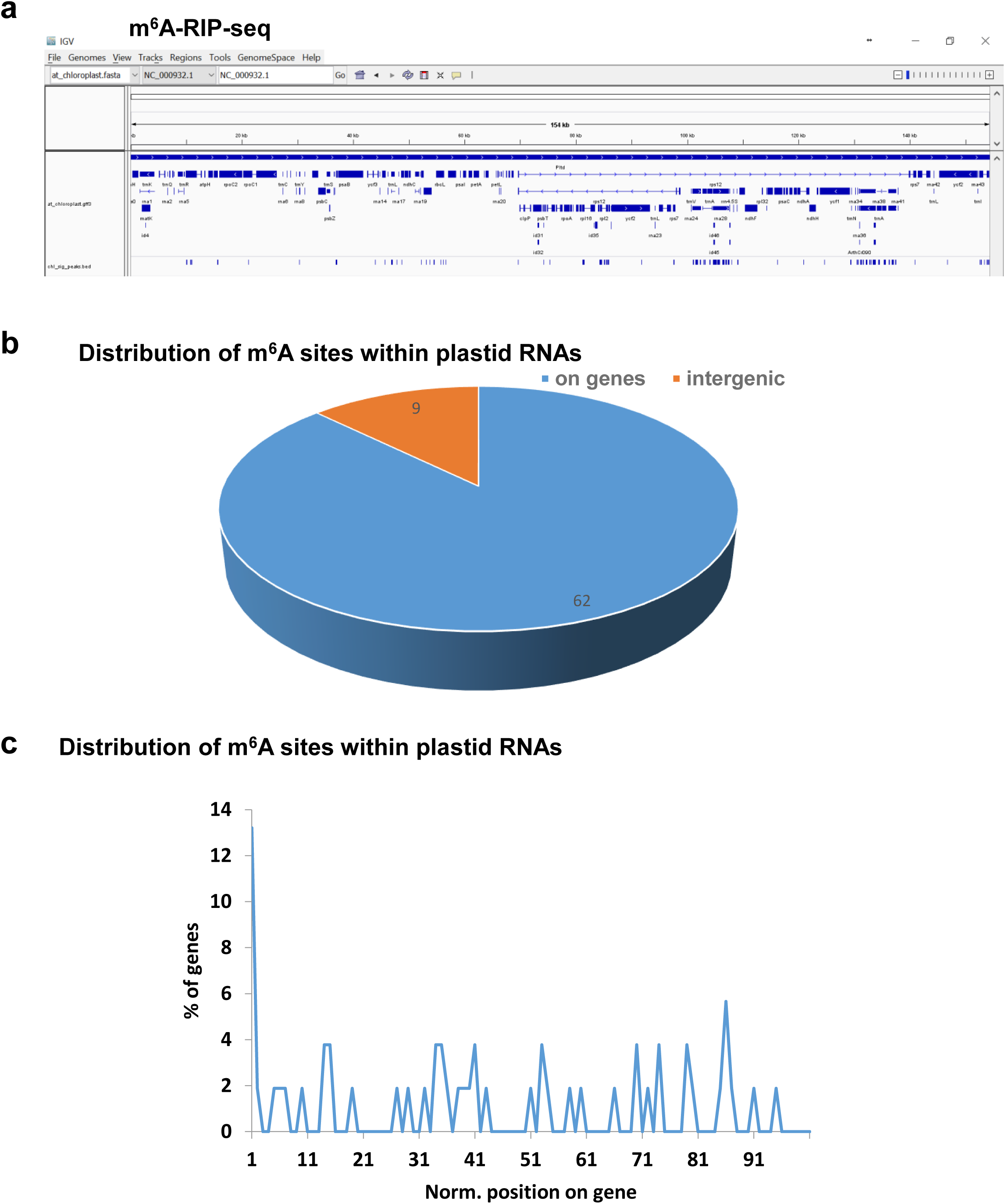
(a) Analysis of m^6^A metylomes in plastid RNAs of Arabidopsis plants. The topology of m^6^A modifications was analyzed by m^6^A-RIP-seq analyses with anti-m^6^A antibodies. The reads were aligned to the chloroplast genome of Arabidopsis (NC_000932.1). (b) pie-charts presenting the fraction of m^6^A peaks in genes (62 reads) and expressed-intergenic regions (9 reads) in Arabidopsis chloroplasts. (c) Distribution of m^6^A peaks in Arabidopsis sequences corresponding to coding regions, according to normalized gene lengths. A list of mitochondrial genes in Arabidopsis and cauliflower identified by the m6A-RIP-seq analyses is presented in Table S6.

**Table S1.** Chromatographic data and MS parameters for m6A nucleoside analysis.

| Nucleoside label | Retention time (min) | Formula | Abundance (relative counts) | Target Mass (gr • mol<sup>-1</sup>) |
| --- | --- | --- | --- | --- |
| Cytidine (C) | 3.69 | C<sub>9</sub>H<sub>13</sub>N<sub>3</sub>O<sub>5</sub> | 12300 | 243.08698 |
| Uridine (U) | 5.20 | C<sub>9</sub>H<sub>12</sub>N<sub>2</sub>O<sub>6</sub> | 1931 | 244.07114 |
| Guanosine (G) | 9.18 | C<sub>10</sub> H<sub>13</sub> N<sub>5</sub> O<sub>5</sub> | 3160 | 283.09308 |
| Adenosine (A) | 12.47 | C<sub>10</sub> H<sub>13</sub> N<sub>5</sub> O<sub>4</sub> | 3113 | 267.09795 |
| *N*<sup>6</sup>-methyladenosine (m<sup>6</sup>A) | 15.79 | C<sub>11</sub> H<sub>15</sub> N<sub>5</sub> O<sub>4</sub> | 2459 | 281.11359 |

**B. ESI-MS and ESI-MS/MS parameters for m<sup>6</sup>A nucleoside analysis**
| Compounds | Precursor ions (m/z) | Product ions (m/z) | Collision energy(V) | ΔT (min) |
| --- | --- | --- | --- | --- |
| M<sup>6</sup>A | 282. 1211[M+H]<sup>+</sup> | 123.06384 [M-159+H]<sup>+</sup> | 15 | 14-17 |
|  |  | 130.99616 [M-151+H]<sup>+</sup> | 15 |  |
|  |  | 133.04954 [M-149+H]<sup>+</sup> | 15 |  |
|  |  | 150.07742 [M-132+H]<sup>+</sup> | 15 |  |

**Table S2.**
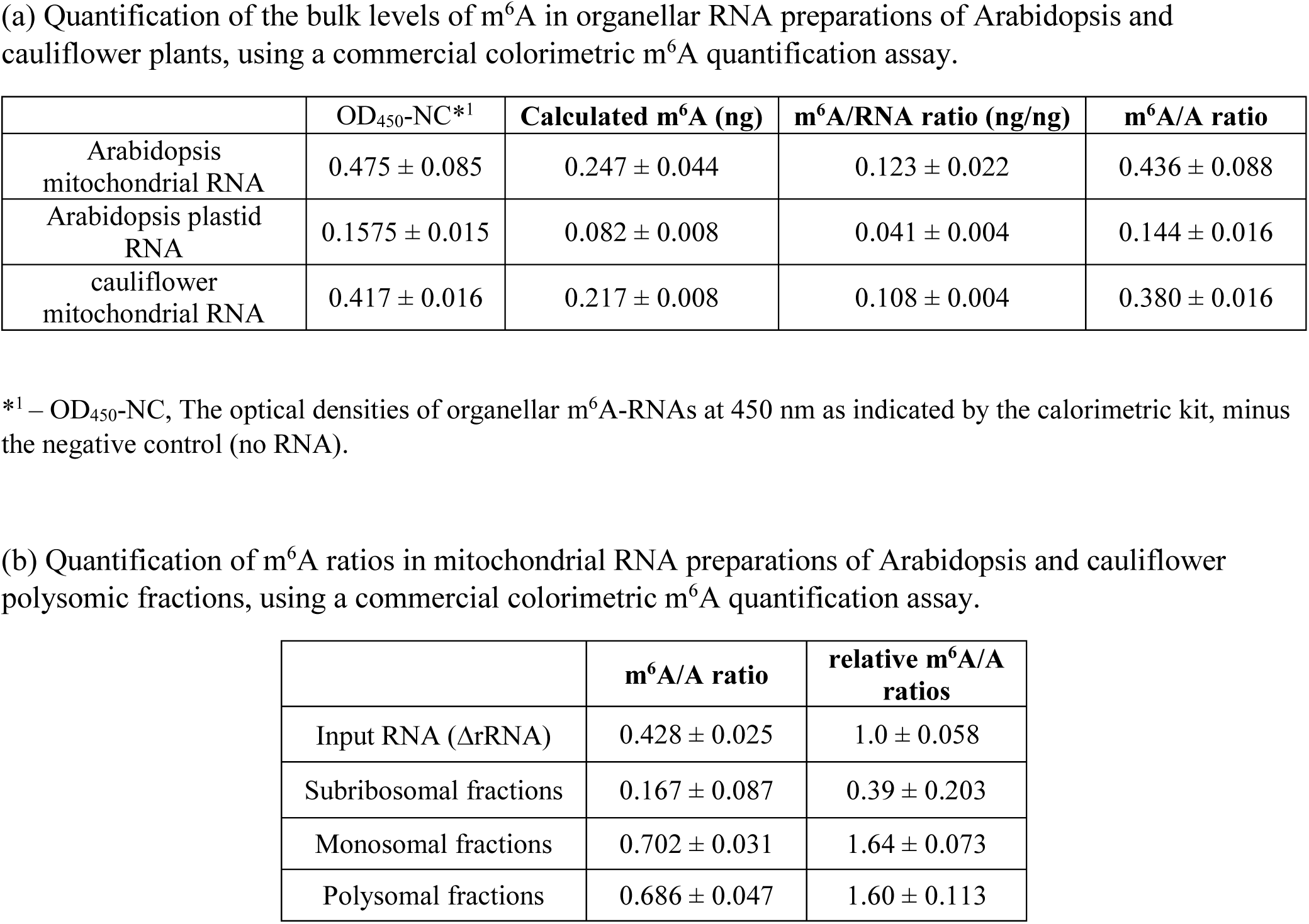
Quantification of the bulk levels of m^6^A by calorimetric assays.

**Table S3.** RNA-seq’s of rRNA-depleted total RNAs from Arabidopsis and cauliflower mitochondria (PRJNA472433, replicate 1).

| Library | Plant | Total reads | QC filtered | Mapped to mtDNA |
| --- | --- | --- | --- | --- |
| Input RNA | *A. thaliana* | 62126071 | 61812498<br>(99.50%) | 25125548<br>(40.65%) |
| m6A-RIP | *A. thaliana* | 7528773 | 7425907<br>(98.63%) | 1327196<br>(17.87%) |
| Input RNA | *B. oleracea* | 51288692 | 51048195<br>(99.53%) | 37467278<br>(73.40%) |
| m6A-RIP | *B. oleracea* | 29371176 | 29201322<br>(99.42%) | 20505049<br>(70.22%) |

**Table S4.**
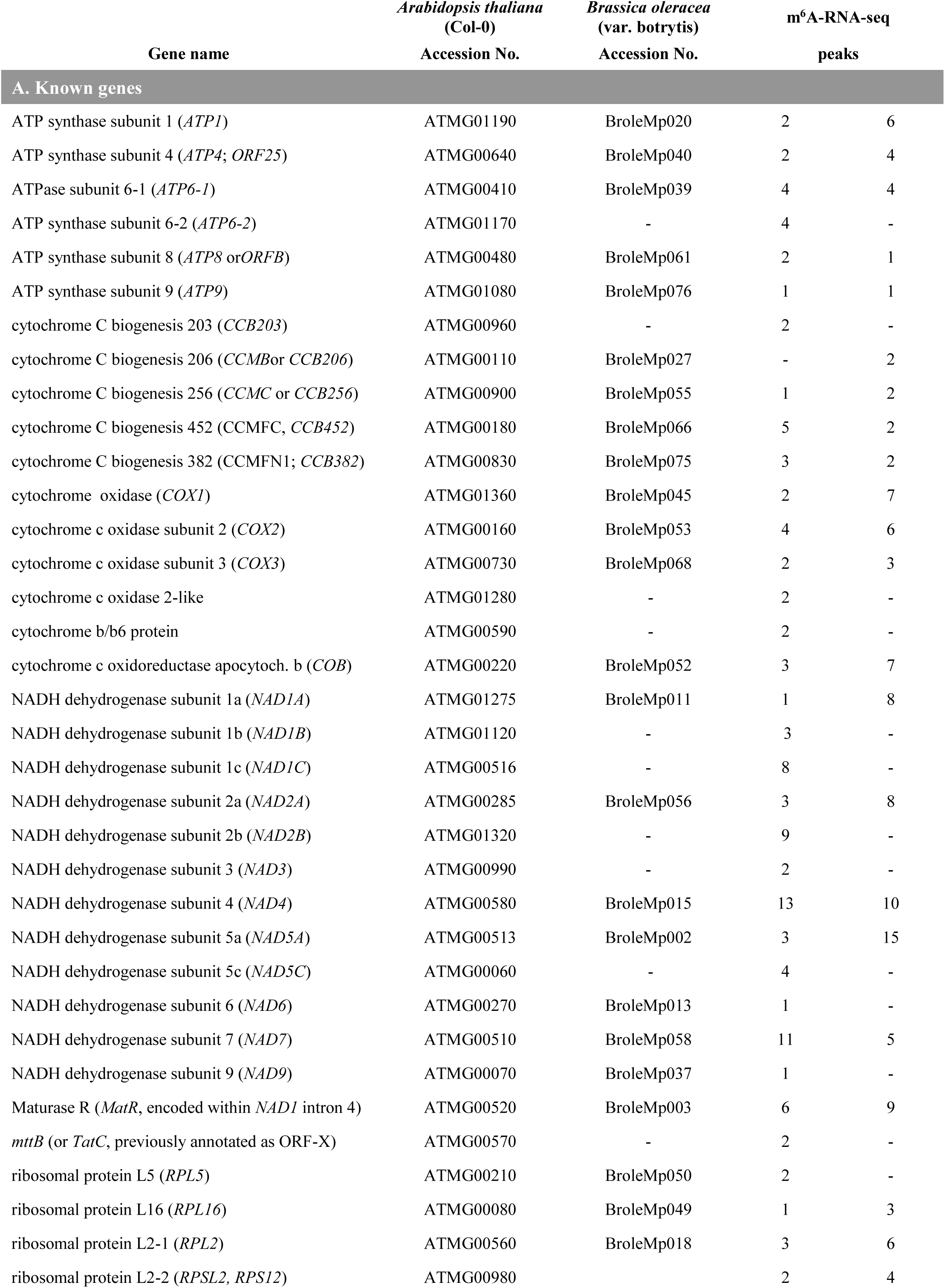
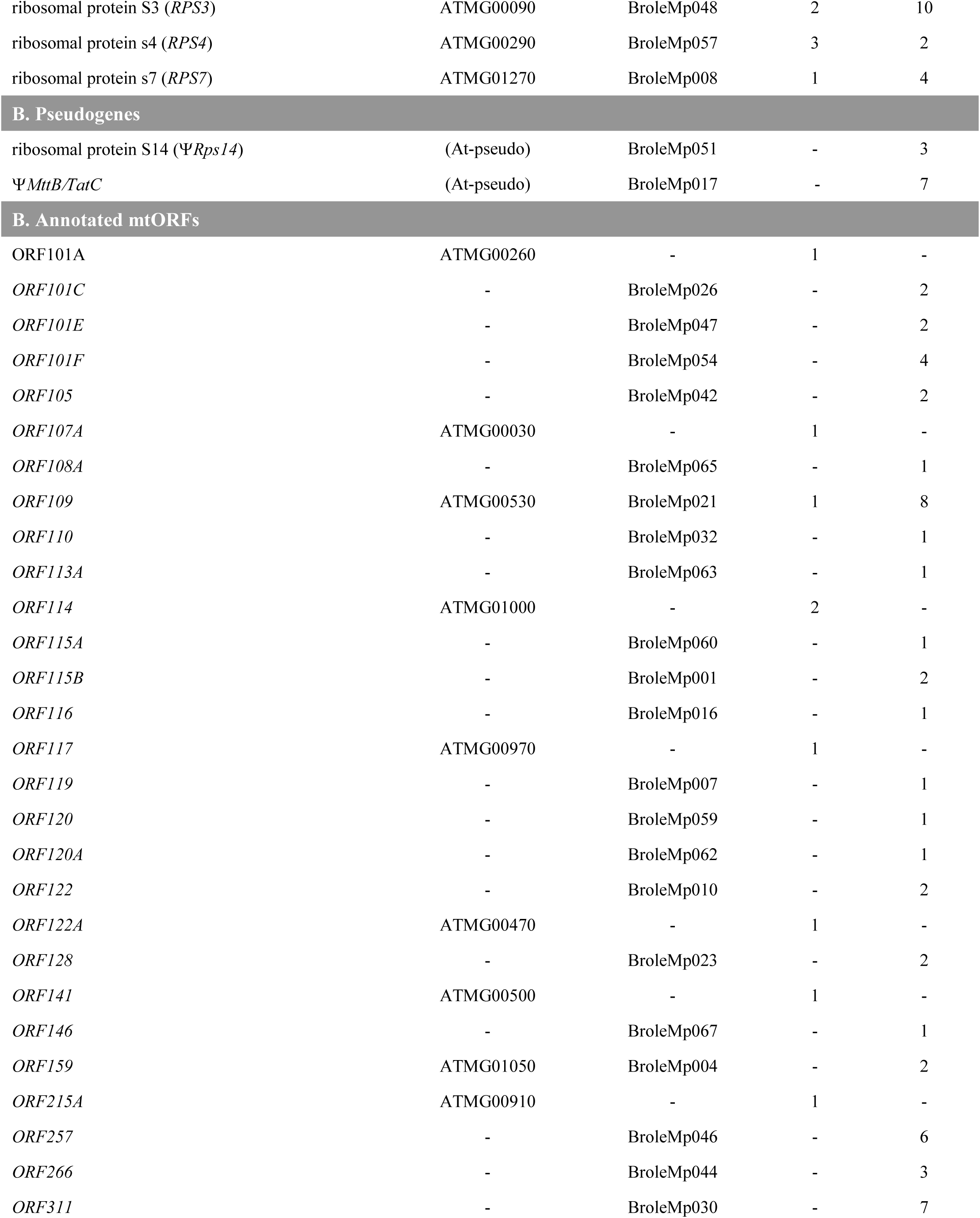
List of mitochondrial genes in Arabidopsis and cauliflower identified by the m6A-RIP-seq analyses (PRJNA472433, replicate 1).

| Gene name | *Arabidopsis thaliana*<br>(Col-0) | *Brassica oleracea*<br>(var. botrytis) | m<sup>6</sup>A-RNA-seq |  |
| --- | --- | --- | --- | --- |
|  | Accession No. | Accession No. | peaks |  |
| A. Known genes |  |  |  |  |
| ATP synthase subunit 1 (*ATP1*) | ATMG01190 | BroleMp020 | 2 | 6 |
| ATP synthase subunit 4 (*ATP4*; *ORF25*) | ATMG00640 | BroleMp040 | 2 | 4 |
| ATPase subunit 6-1 (*ATP6-1*) | ATMG00410 | BroleMp039 | 4 | 4 |
| ATP synthase subunit 6-2 (*ATP6-2*) | ATMG01170 | - | 4 | - |
| ATP synthase subunit 8 (*ATP8* or*ORFB*) | ATMG00480 | BroleMp061 | 2 | 1 |
| ATP synthase subunit 9 (*ATP9*) | ATMG01080 | BroleMp076 | 1 | 1 |
| cytochrome C biogenesis 203 (*CCB203*) | ATMG00960 | - | 2 | - |
| cytochrome C biogenesis 206 (*CCMB*or *CCB206*) | ATMG00110 | BroleMp027 | - | 2 |
| cytochrome C biogenesis 256 (*CCMC* or *CCB256*) | ATMG00900 | BroleMp055 | 1 | 2 |
| cytochrome C biogenesis 452 (*CCMFC*, *CCB452*) | ATMG00180 | BroleMp066 | 5 | 2 |
| cytochrome C biogenesis 382 (*CCMFN1*; *CCB382*) | ATMG00830 | BroleMp075 | 3 | 2 |
| cytochrome oxidase (*COX1*) | ATMG01360 | BroleMp045 | 2 | 7 |
| cytochrome c oxidase subunit 2 (*COX2*) | ATMG00160 | BroleMp053 | 4 | 6 |
| cytochrome c oxidase subunit 3 (*COX3*) | ATMG00730 | BroleMp068 | 2 | 3 |
| cytochrome c oxidase 2-like | ATMG01280 | - | 2 | - |
| cytochrome b/b6 protein | ATMG00590 | - | 2 | - |
| cytochrome c oxidoreductase apocytoch. b (*COB*) | ATMG00220 | BroleMp052 | 3 | 7 |
| NADH dehydrogenase subunit 1a (*NAD1A*) | ATMG01275 | BroleMp011 | 1 | 8 |
| NADH dehydrogenase subunit 1b (*NAD1B*) | ATMG01120 | - | 3 | - |
| NADH dehydrogenase subunit 1c (*NAD1C*) | ATMG00516 | - | 8 | - |
| NADH dehydrogenase subunit 2a (*NAD2A*) | ATMG00285 | BroleMp056 | 3 | 8 |
| NADH dehydrogenase subunit 2b (*NAD2B*) | ATMG01320 | - | 9 | - |
| NADH dehydrogenase subunit 3 (*NAD3*) | ATMG00990 | - | 2 | - |
| NADH dehydrogenase subunit 4 (*NAD4*) | ATMG00580 | BroleMp015 | 13 | 10 |
| NADH dehydrogenase subunit 5a (*NAD5A*) | ATMG00513 | BroleMp002 | 3 | 15 |
| NADH dehydrogenase subunit 5c (*NAD5C*) | ATMG00060 | - | 4 | - |
| NADH dehydrogenase subunit 6 (*NAD6*) | ATMG00270 | BroleMp013 | 1 | - |
| NADH dehydrogenase subunit 7 (*NAD7*) | ATMG00510 | BroleMp058 | 11 | 5 |
| NADH dehydrogenase subunit 9 (*NAD9*) | ATMG00070 | BroleMp037 | 1 | - |
| Maturase R (*MatR*, encoded within *NAD1* intron 4) | ATMG00520 | BroleMp003 | 6 | 9 |
| *mttB* (or *TatC*, previously annotated as ORF-X) | ATMG00570 | - | 2 | - |
| ribosomal protein L5 (*RPL5*) | ATMG00210 | BroleMp050 | 2 | - |
| ribosomal protein L16 (*RPL16*) | ATMG00080 | BroleMp049 | 1 | 3 |
| ribosomal protein L2-1 (*RPL2*) | ATMG00560 | BroleMp018 | 3 | 6 |
| ribosomal protein L2-2 (*RPSL2*, *RPS12*) | ATMG00980 |  | 2 | 4 |
| ribosomal protein S3 (*RPS3*) | ATMG00090 | BroleMp048 | 2 | 10 |
| ribosomal protein s4 (*RPS4*) | ATMG00290 | BroleMp057 | 3 | 2 |
| ribosomal protein s7 (*RPS7*) | ATMG01270 | BroleMp008 | 1 | 4 |
| B. Pseudogenes |  |  |  |  |
| ribosomal protein S14 (*ΨRps14*) | (At-pseudo) | BroleMp051 | - | 3 |
| *ΨMttB/TatC* | (At-pseudo) | BroleMp017 | - | 7 |
| B. Annotated mtORFs |  |  |  |  |
| ORF101A | ATMG00260 | - | 1 | - |
| *ORF101C* | - | BroleMp026 | - | 2 |
| *ORF101E* | - | BroleMp047 | - | 2 |
| *ORF101F* | - | BroleMp054 | - | 4 |
| *ORF105* | - | BroleMp042 | - | 2 |
| *ORF107A* | ATMG00030 | - | 1 | - |
| *ORF108A* | - | BroleMp065 | - | 1 |
| *ORF109* | ATMG00530 | BroleMp021 | 1 | 8 |
| *ORF110* | - | BroleMp032 | - | 1 |
| *ORF113A* | - | BroleMp063 | - | 1 |
| *ORF114* | ATMG01000 | - | 2 | - |
| *ORF115A* | - | BroleMp060 | - | 1 |
| *ORF115B* | - | BroleMp001 | - | 2 |
| *ORF116* | - | BroleMp016 | - | 1 |
| *ORF117* | ATMG00970 | - | 1 | - |
| *ORF119* | - | BroleMp007 | - | 1 |
| *ORF120* | - | BroleMp059 | - | 1 |
| *ORF120A* | - | BroleMp062 | - | 1 |
| *ORF122* | - | BroleMp010 | - | 2 |
| *ORF122A* | ATMG00470 | - | 1 | - |
| *ORF128* | - | BroleMp023 | - | 2 |
| *ORF141* | ATMG00500 | - | 1 | - |
| *ORF146* | - | BroleMp067 | - | 1 |
| *ORF159* | ATMG01050 | BroleMp004 | - | 2 |
| *ORF215A* | ATMG00910 | - | 1 | - |
| *ORF257* | - | BroleMp046 | - | 6 |
| *ORF266* | - | BroleMp044 | - | 3 |
| *ORF311* | - | BroleMp030 | - | 7 |

**Table S5.**
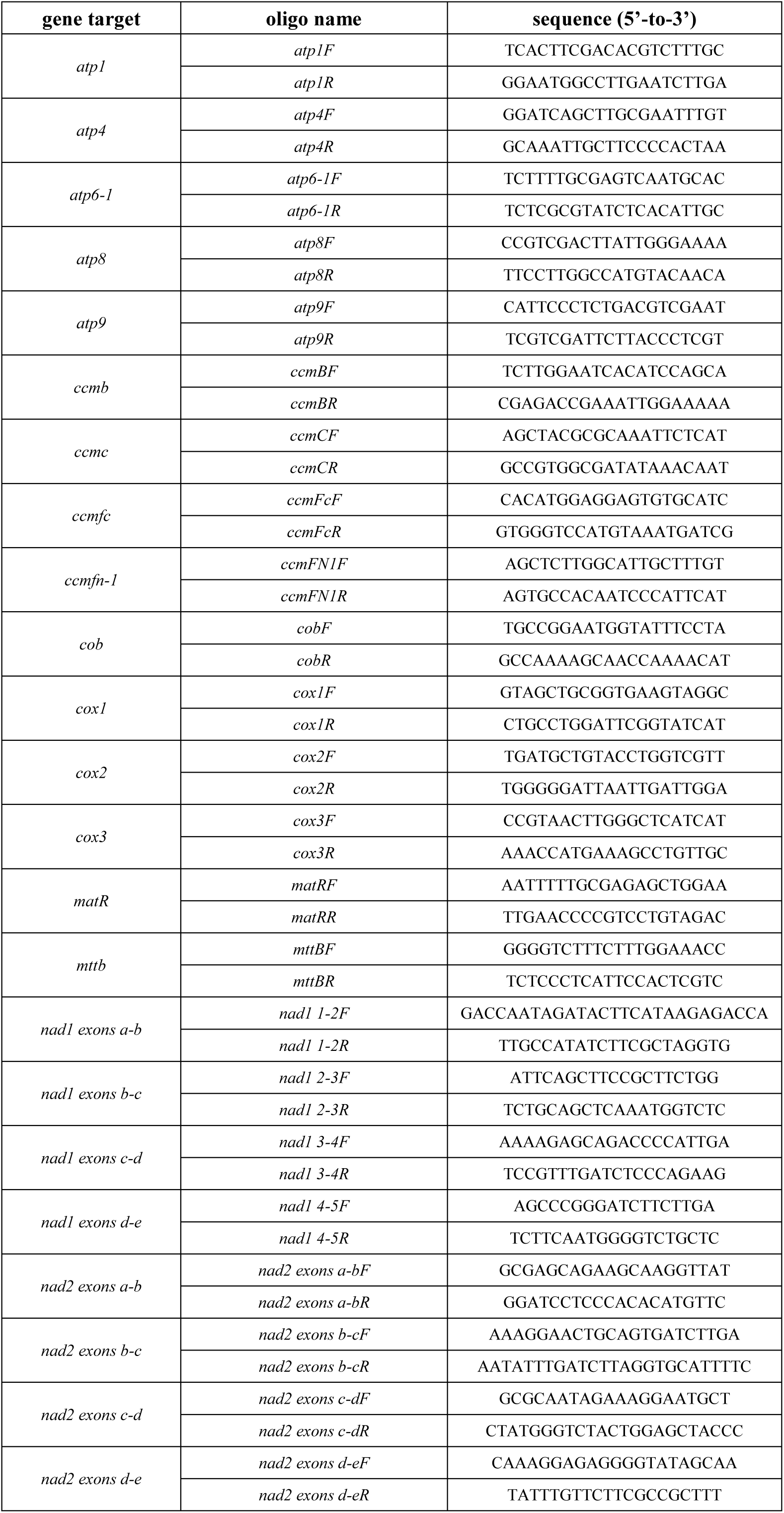
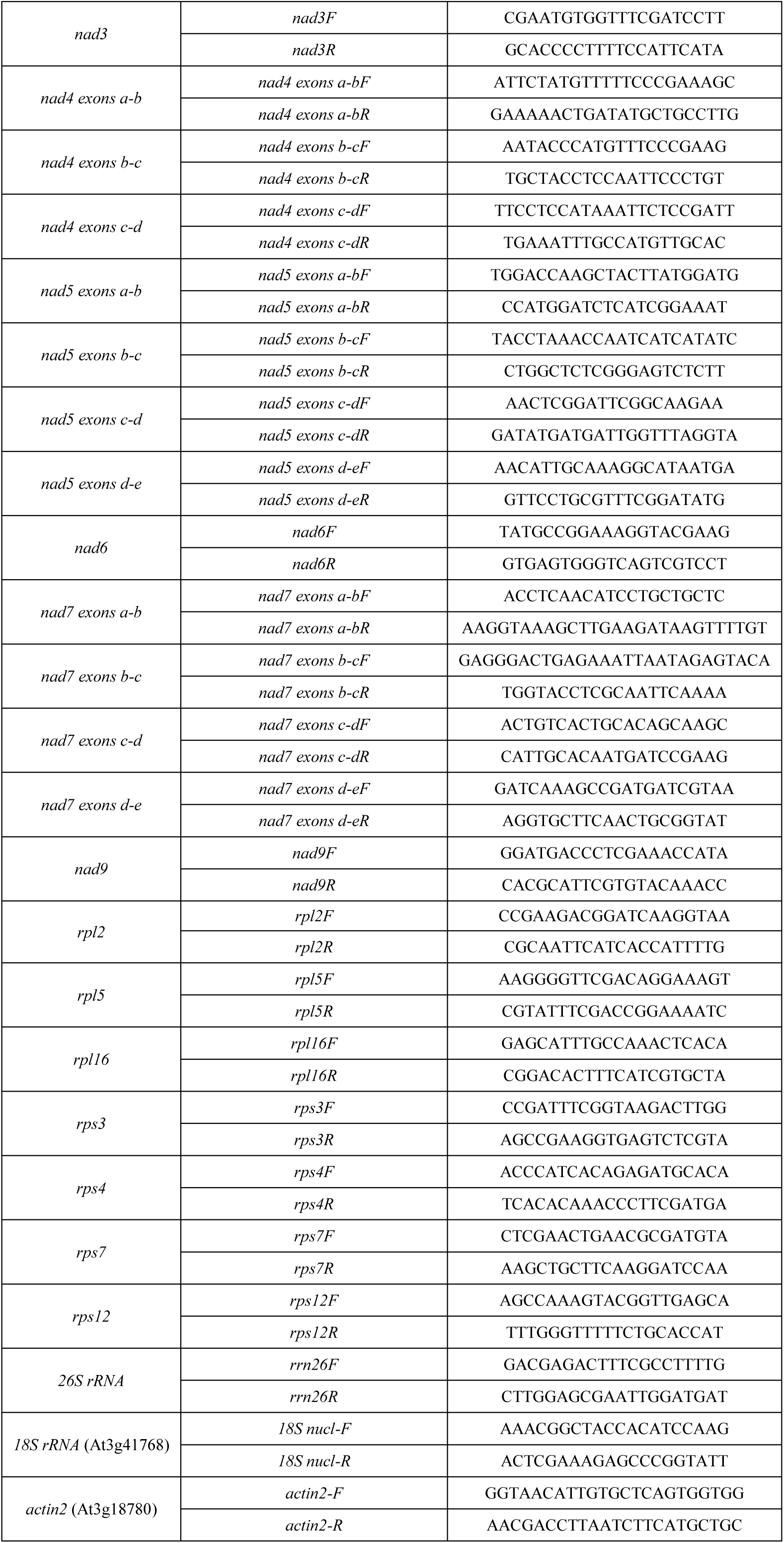
Lists of oligonucleotides used for the analysis of the splicing profiles of wild-type and mutant plants by RT-qPCR experiments.

**Table S6.**
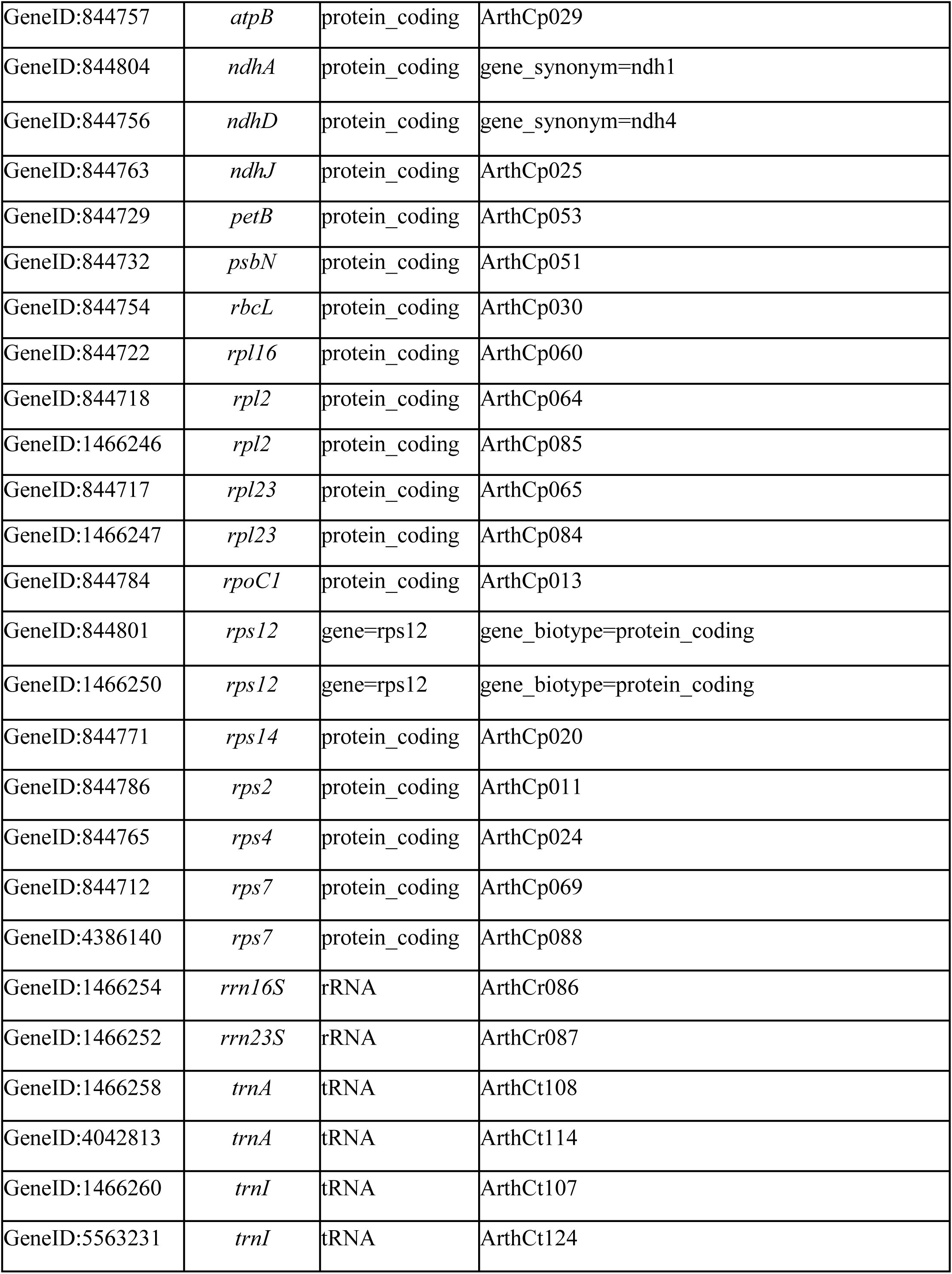
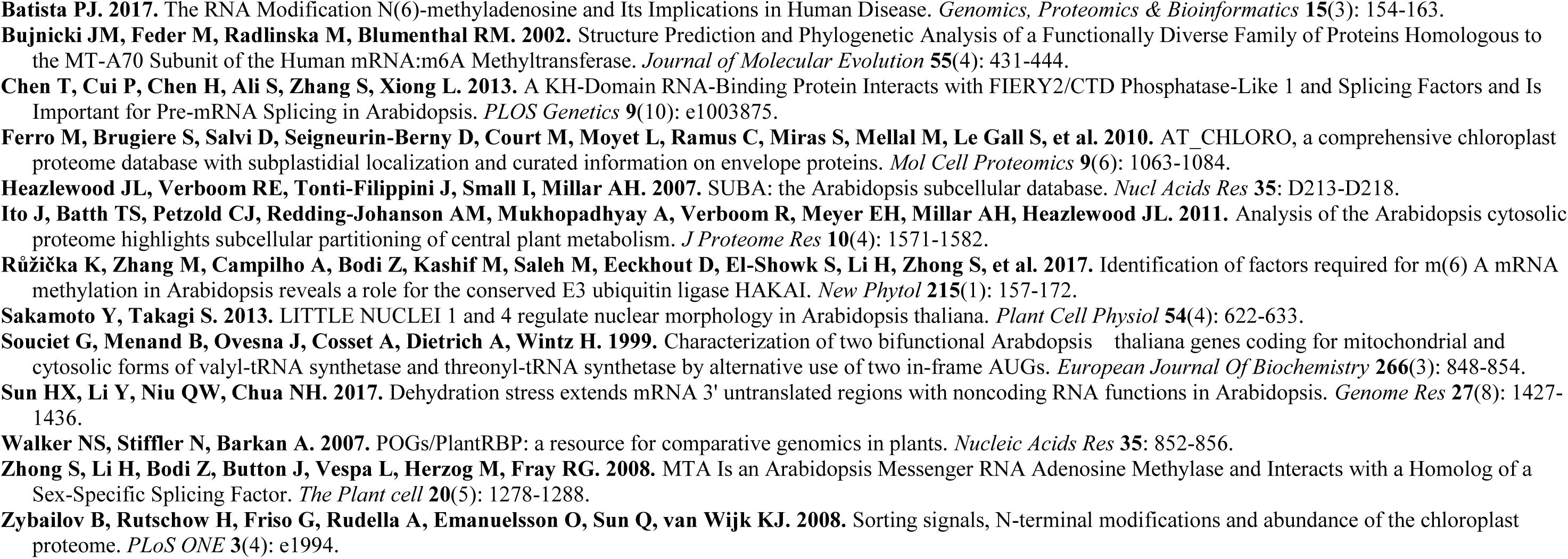
List of chloroplast genes in Arabidopsis identified by the m^6^A-RIP-seq analyses.

**Table S7.**
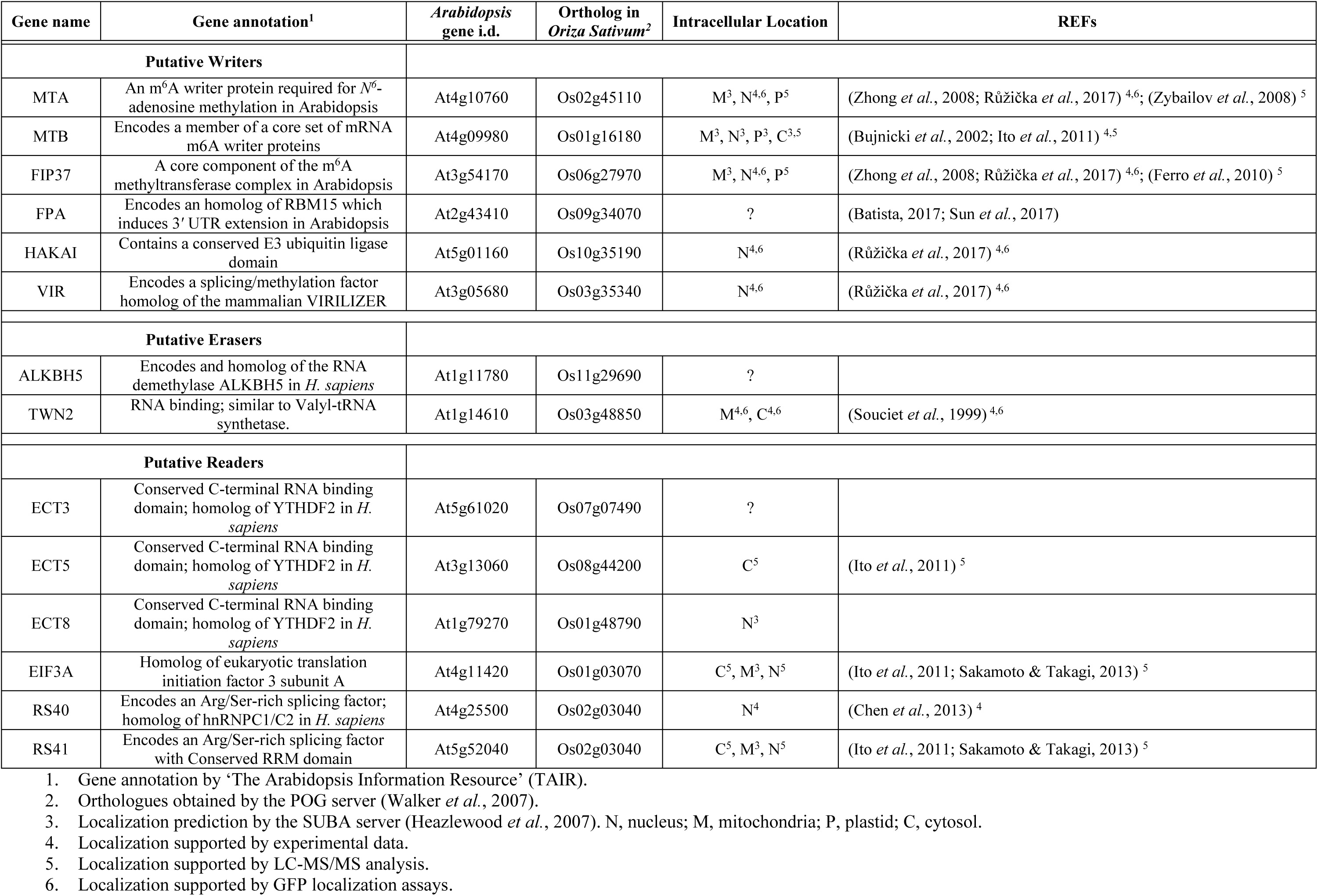
List of putative m6A modification enzymes in Arabidopsis mitochondria.

**Table S8.**
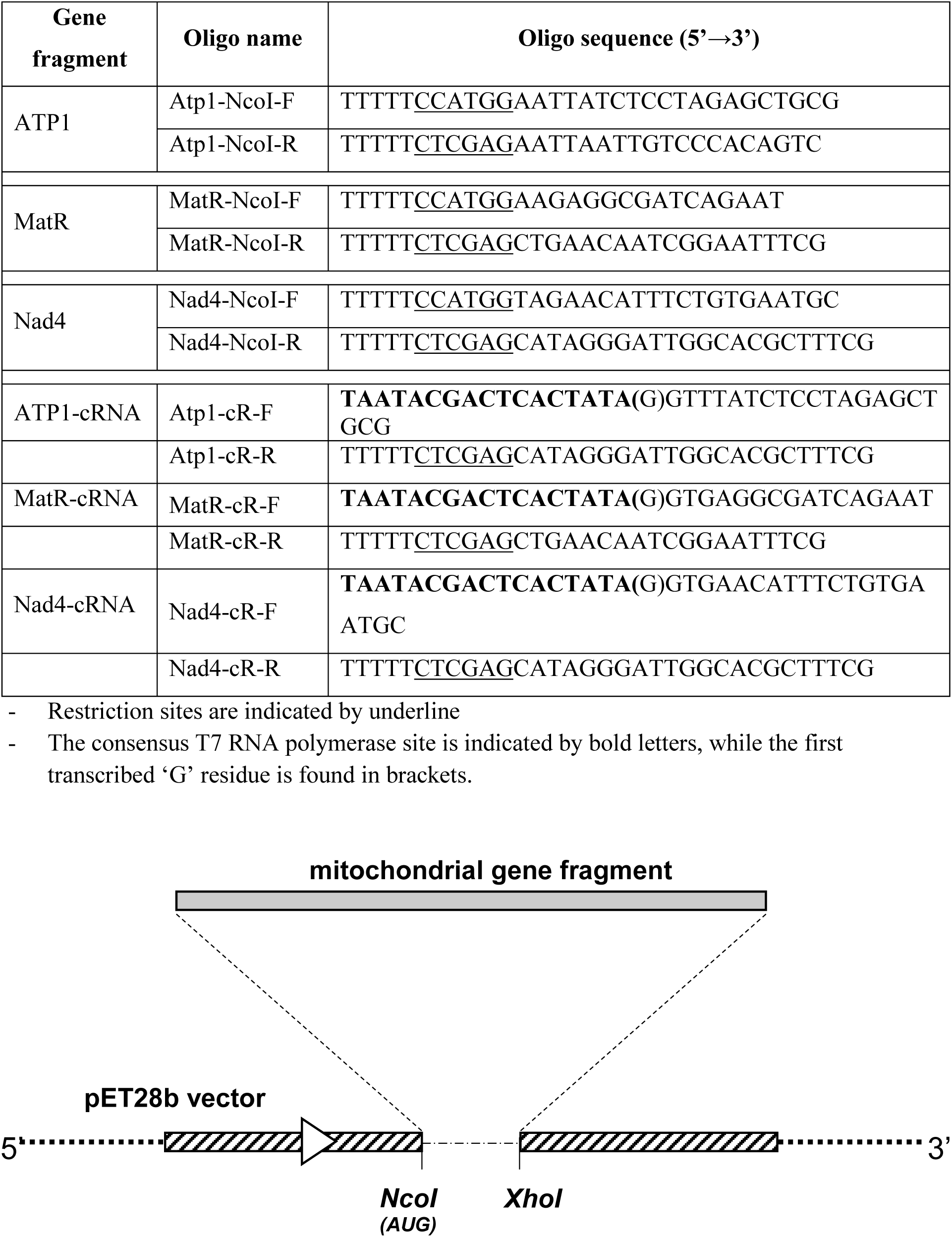
Oligonucleotides used to the construction of gene-fragments corresponding to MatR, NAD4 and ATP1.

## References

Akichika, S., Hirano, S., Shichino, Y., Suzuki, T., Nishimasu, H., Ishitani, R., Sugita, A., Hirose, Y., Iwasaki, S., Nureki, O. and Suzuki, T. (2019) Cap-specific terminal N (6)-methylation of RNA by an RNA polymerase II-associated methyltransferase. Science, 363.

Bailey, T.L., Boden, M., Buske, F.A., Frith, M., Grant, C.E., Clementi, L., Ren, J., Li, W.W. and Noble, W.S. (2009) MEME SUITE: tools for motif discovery and searching. Nucleic Acids Res, 37, W202–208.

Barkan, A. (1998) Approaches to investigating nuclear genes that function in chloroplast biogenesis in land plants. In Methods in Enzymology (Lee, M. ed: Academic Press, pp. 38–57.

Barkan, A., Klipcan, L., Ostersetzer, O., Kawamura, T., Asakura, Y. and Watkins, K.P. (2007) The CRM domain: An RNA binding module derived from an ancient ribosome-associated protein. RNA, 13, 55–64.

Barkan, A. and Small, I. (2014) Pentatricopeptide Repeat Proteins in Plants. Annual Review of Plant Biology, 65, 415–442.

Binder, S., Stoll, K. and Stoll, B. (2016) Maturation of 5’ ends of plant mitochondrial RNAs. Physiol Plant.

Bodi, Z., Zhong, S., Mehra, S., Song, J., Graham, N., Li, H., May, S. and Fray, R.G. (2012) Adenosine Methylation in Arabidopsis mRNA is Associated with the 3’ End and Reduced Levels Cause Developmental Defects. Front Plant Sci, 3, 48.

Bogsch, E.G., Sargent, F., Stanley, N.R., Berks, B.C., Robinson, C. and Palmer, T. (1998) AN essential component of a novel bacterial protein export system with homologues in plastids and mitochondria. J. Biol. Chem., 273, 18003–18006.

Bolger, A.M., Lohse, M. and Usadel, B. (2014) Trimmomatic: a flexible trimmer for Illumina sequence data. Bioinformatics, 30, 2114–2120.

Bonen, L. (2018) Mitochondrial Genomes in Land Plants, pp. 734–742.

Burgess, A., David, R. and Searle, I.R. (2016) Deciphering the epitranscriptome: A green perspective. J Integr Plant Biol, 58, 822–835.

Cermakian, N., Ikeda, T.M., Cedergren, R. and Gray, M.W. (1996) Sequences homologous to yeast mitochondrial and bacteriophage T3 and T7 RNA polymerases are widespread throughout the eukaryotic lineage. Nucleic Acids Res, 24, 648–654.

Chateigner-Boutin, A.-L. and Small, I. (2010) Plant RNA editing. RNA biology, 7, 213–219.

Chen, L. and Liu, Y.G. (2014) Male sterility and fertility restoration in crops. Annu Rev Plant Biol, 65, 579–606.

Chen, M., Urs, M.J., Sánchez-González, I., Olayioye, M.A., Herde, M. and Witte, C.-P. (2018) m6A RNA Degradation Products Are Catabolized by an Evolutionarily Conserved N6-Methyl-AMP Deaminase in Plant and Mammalian Cells. The Plant Cell, 30, 1511–1522.

Choi, J., Ieong, K.W., Demirci, H., Chen, J., Petrov, A., Prabhakar, A., O’Leary, S.E., Dominissini, D., Rechavi, G., Soltis, S.M., Ehrenberg, M. and Puglisi, J.D. (2016) N(6)-methyladenosine in mRNA disrupts tRNA selection and translation-elongation dynamics. Nat Struct Mol Biol, 23, 110–115.

Cohen, S., Zmudjak, M., Colas des Francs-Small, C., Malik, S., Shaya, F., Keren, I., Belausov, E., Many, Y., Brown, G.G., Small, I. and Ostersetzer-Biran, O. (2014) nMAT4, a maturase factor required for nad1 pre-mRNA processing and maturation, is essential for holocomplex I biogenesis in Arabidopsis mitochondria. The Plant Journal, 78, 253–268.

Colas des Francs-Small, C. and Small, I. (2014) Surrogate mutants for studying mitochondrially encoded functions. Biochimie, 100, 234–242.

Dai, Q., Fong, R., Saikia, M., Stephenson, D., Yu, Y.-t., Pan, T. and Piccirilli, J.A. (2007) Identification of recognition residues for ligation-based detection and quantitation of pseudouridine and N6 -methyladenosine. Nucleic Acids Research, 35, 6322–6329.

Deng, X., Chen, K., Luo, G.Z., Weng, X., Ji, Q., Zhou, T. and He, C. (2015) Widespread occurrence of N6-methyladenosine in bacterial mRNA. Nucleic Acids Res, 43, 6557–6567.

Desrosiers, R., Friderici, K. and Rottman, F. (1974) Identification of methylated nucleosides in messenger RNA from Novikoff hepatoma cells. Proc Natl Acad Sci U S A, 71, 3971–3975.

Dominissini, D., Moshitch-Moshkovitz, S., Schwartz, S., Salmon-Divon, M., Ungar, L., Osenberg, S., Cesarkas, K., Jacob-Hirsch, J., Amariglio, N., Kupiec, M., Sorek, R. and Rechavi, G. (2012) Topology of the human and mouse m6A RNA methylomes revealed by m6A-seq. Nature, 485, 201–206.

Engel, M., Roeh, S., Eggert, C., Kaplick, P.M., Tietze, L., Arloth, J., Weber, P., Rex-Haffner, M., Jakovcevski, M., Uhr, M., Eder, M., Wotjak, C.T., Schmidt, M.V., Deussing, J.M., Binder, E.B. and Chen, A. (2017) The role of m6A-RNA methylation in stress response regulation. bioRxiv, 1–81.

Fernie, A.R., Carrari, F. and Sweetlove, L.J. (2004) Respiratory metabolism: glycolysis, the TCA cycle and mitochondrial electron transport. Curr Opin Plant Biol, 7, 254–261.

Ge, J. and Yu, Y.-T. (2013) RNA pseudouridylation: new insights into an old modification. Trends in biochemical sciences, 38, 210–218.

Grewe, F., Edger, P.P., Keren, I., Sultan, L., Pires, J.C., Ostersetzer-Biran, O. and Mower, J.P. (2014) Comparative analysis of 11 Brassicales mitochondrial genomes and the mitochondrial transcriptome of Brassica oleracea. Mitochondrion, 19, Part B, 135–143.

Gualberto, J.M., Mileshina, D., Wallet, C., Niazi, A.K., Weber-Lotfi, F. and Dietrich, A. (2014) The plant mitochondrial genome: Dynamics and maintenance. Biochimie, 100, 107–120.

Gualberto, J.M. and Newton, K.J. (2017) Plant Mitochondrial Genomes: Dynamics and Mechanisms of Mutation. Annu Rev Plant Biol, 68, 225–252.

Guo, W., Grewe, F., Fan, W., Young, G.J., Knoop, V., Palmer, J.D. and Mower, J.P. (2016) Ginkgo and Welwitschia Mitogenomes Reveal Extreme Contrasts in Gymnosperm Mitochondrial Evolution. Mol Biol Evol.

Hammani, K. and Giege, P. (2014) RNA metabolism in plant mitochondria. Trends Plant Sci, 19, 380–389.

Heazlewood, J.L., Whelan, J. and Millar, A.H. (2003) The products of the mitochondrial orf25 and orfB genes are FO components in the plant F1FO ATP synthase. FEBS Lett, 540, 201–205.

Heinz, S., Benner, C., Spann, N., Bertolino, E., Lin, Y.C., Laslo, P., Cheng, J.X., Murre, C., Singh, H. and Glass, C.K. (2010) Simple combinations of lineage-determining transcription factors prime cis-regulatory elements required for macrophage and B cell identities. Mol Cell, 38, 576–589.

Hoernes, T.P., Clementi, N., Faserl, K., Glasner, H., Breuker, K., Lindner, H., Huttenhofer, A. and Erlacher, M.D. (2016) Nucleotide modifications within bacterial messenger RNAs regulate their translation and are able to rewire the genetic code. Nucleic Acids Res, 44, 852–862.

Ichinose, M. and Sugita, M. (2016a) RNA Editing and Its Molecular Mechanism in Plant Organelles. Genes, 8, 5–11.

Ichinose, M. and Sugita, M. (2016b) RNA Editing and Its Molecular Mechanism in Plant Organelles. Genes, 8.

John, U., Lu, Y., Wohlrab, S., Groth, M., Janouškovec, J., Kohli, G.S., Mark, F.C., Bickmeyer, U., Farhat, S., Felder, M., Frickenhaus, S., Guillou, L., Keeling, P.J., Moustafa, A., Porcel, B.M., Valentin, K. and Glöckner, G. (2019) An aerobic eukaryotic parasite with functional mitochondria that likely lacks a mitochondrial genome. Science Advances, 5, eaav1110.

Kahlau, S. and Bock, R. (2008) Plastid transcriptomics and translatomics of tomato fruit development and chloroplast-to-chromoplast differentiation: Chromoplast gene expression largely serves the production of a single protein. Plant Cell, 20, 856–874.

Ke, S., Alemu, E.A., Mertens, C., Gantman, E.C., Fak, J.J., Mele, A., Haripal, B., Zucker-Scharff, I., Moore, M.J., Park, C.Y., Vågbø, C.B., Kusśnierczyk, A., Klungland, A., Darnell, J.E. and Darnell, R.B. (2015) A majority of m6A residues are in the last exons, allowing the potential for 3′ UTR regulation. Genes & Development, 29, 2037–2053.

Keren, I., Bezawork-Geleta, A., Kolton, M., Maayan, I., Belausov, E., Levy, M., Mett, A., Gidoni, D., Shaya, F. and Ostersetzer-Biran, O. (2009) AtnMat2, a nuclear-encoded maturase required for splicing of group-II introns in Arabidopsis mitochondria. RNA, 15, 2299–2311.

Keren, I., Klipcan, L., Bezawork-Geleta, A., Kolton, M., Shaya, F. and Ostersetzer-Biran, O. (2008) Characterization of the molecular basis of group II intron RNA recognition by CRS1-CRM domains. J Biol Chem, 283, 23333–23342.

Keren, I., Shaya, F. and Ostersetzer-Biran, O. (2011) An optimized method for the analysis of plant mitochondria RNAs by northern-blotting. Endocy Cell Res, 1, 34–42.

Keren, I., Tal, L., Colas des Francs-Small, C., Araújo, W.L., Shevtsov, S., Shaya, F., Fernie, A.R., Small, I. and Ostersetzer-Biran, O. (2012) nMAT1, a nuclear-encoded maturase involved in the *trans*-splicing of *nad1* intron 1, is essential for mitochondrial complex I assembly and function. Plant J, 71, 413–426.

Knoop, V. (2012) Seed Plant Mitochondrial Genomes: Complexity Evolving. In Genomics of chloroplasts and mitochondria (Bock, R. and Knoop, V. eds), pp. 175–200.

Langmead, B. and Salzberg, S.L. (2012) Fast gapped-read alignment with Bowtie 2. Nature methods, 9, 357–359.

Li, Y., Wang, X., Li, C., Hu, S., Yu, J. and Song, S. (2014) Transcriptome-wide N(6)-methyladenosine profiling of rice callus and leaf reveals the presence of tissue-specific competitors involved in selective mRNA modification. RNA biology, 11, 1180–1188.

Liere, K., Weihe, A. and Börner, T. (2011) The transcription machineries of plant mitochondria and chloroplasts: Composition, function, and regulation. Journal of Plant Physiology, 168, 1345–1360.

Liu, N., Dai, Q., Zheng, G., He, C., Parisien, M. and Pan, T. (2015) N(6)-methyladenosine-dependent RNA structural switches regulate RNA-protein interactions. Nature, 518, 560–564.

Liu, N. and Pan, T. (2016) N6-methyladenosine-encoded epitranscriptomics. Nat Struct Mol Biol, 23, 98–102.

Luo, G.-Z., MacQueen, A., Zheng, G., Duan, H., Dore, L.C., Lu, Z., Liu, J., Chen, K., Jia, G., Bergelson, J. and He, C. (2014) Unique features of the m6A methylome in Arabidopsis thaliana. Nature communications, 5, 5630.

Maclean, A.E., Hertle, A.P., Ligas, J., Bock, R., Balk, J. and Meyer, E.H. (2018) Absence of Complex I Is Associated with Diminished Respiratory Chain Function in European Mistletoe. Curr Biol, 28, 1614–1619.e1613.

Maity, A. and Das, B. (2016) N6-methyladenosine modification in mRNA: machinery, function and implications for health and diseases. Febs J, 283, 1607–1630.

Masters, B.S., Stohl, L.L. and Clayton, D.A. (1987) Yeast mitochondrial RNA polymerase is homologous to those encoded by bacteriophages T3 and T7. Cell, 51, 89–99.

Meyer, K.D. and Jaffrey, S.R. (2017) Rethinking m6A Readers, Writers, and Erasers. Annual Review of Cell and Developmental Biology, 33, 319–342.

Meyer, Kate D., Patil, Deepak P., Zhou, J., Zinoviev, A., Skabkin, Maxim A., Elemento, O., Pestova, Tatyana V., Qian, S.-B. and Jaffrey, Samie R. (2015) 5′ UTR m6A Promotes Cap-Independent Translation. Cell, 163, 999–1010.

Millar, A.H., Whelan, J., Soole, K.L. and Day, D.A. (2011) Organization and regulation of mitochondrial respiration in plants. Ann Rev Plant Biol, 62, 79–104.

Moller, I.M. (2001) PLANT MITOCHONDRIA AND OXIDATIVE STRESS: Electron Transport, NADPH Turnover, and Metabolism of Reactive Oxygen Species. Annu Rev Plant Physiol Plant Mol Biol, 52, 561–591.

Mower, J.P., Sloan, D.B. and Alverson, A.J. (2012) Plant mitochondrial genome diversity: the genomics revolution. In: Plant Genome Diversity Volume 1: Plant Genomes, their Residents, and their Evolutionary Dynamics. Wendel JF, *Greilhuber* *J*, Dolezel J, Leitch IJ (eds). Springer:Dordrecht. pp 123–144.

Nachtergaele, S. and He, C. (2017) The emerging biology of RNA post-transcriptional modifications. RNA biology, 14, 156–163.

Nakagawa, N. and Sakurai, N. (2006) A mutation in At-nMat1a, which encodes a nuclear gene having high similarity to group II Intron maturase, causes impaired splicing of mitochondrial nad4 transcript and altered carbon metabolism in Arabidopsis thaliana. Plant Cell Physiol, 47, 772–783.

Neuwirt, J., Takenaka, M., Van Der Merwe, J.A. and Brennicke, A. (2005) An in vitro RNA editing system from cauliflower mitochondria: Editing site recognition parameters can vary in different plant species. RNA, 11, 1563–1570.

Niu, Y., Zhao, X., Wu, Y.-S., Li, M.-M., Wang, X.-J. and Yang, Y.-G. (2013) N(6)-methyl-adenosine (m(6)A) in RNA: An Old Modification with A Novel Epigenetic Function. *Genomics*, Proteomics & Bioinformatics, 11, 8–17.

Ostersetzer, O. and Adam, Z. (1997) Light-stimulated degradation of an unassembled Rieske FeS protein by a thylakoid-bound protease: The possible role of the FtsH protease. Plant Cell, 9, 957–965.

Ostersetzer, O. and Adam, Z. (1999) Construction of a vector for coupled in vitro transcription/translation. Biotechniques, 27, 428-+.

Ostersetzer, O., Cooke, A.M., Watkins, K.P. and Barkan, A. (2005) CRS1, a Chloroplast Group II Intron Splicing Factor, Promotes Intron Folding through Specific Interactions with Two Intron Domains. The Plant Cell, 17, 241–255.

Park, S., Grewe, F., Zhu, A., Ruhlman, T.A., Sabir, J., Mower, J.P. and Jansen, R.K. (2015) Dynamic evolution of Geranium mitochondrial genomes through multiple horizontal and intracellular gene transfers. New Phytol, 208, 570–583.

Parker, M.T., Knop, K., Sherwood, A.V., Schurch, N.J., Mackinnon, K., Gould, P.D., Hall, A., Barton, G.J. and Simpson, G.G. (2019) Nanopore direct RNA sequencing maps an Arabidopsis N6 methyladenosine epitranscriptome. bioRxiv, 706002.

Patil, D.P., Pickering, B.F. and Jaffrey, S.R. (2018) Reading m(6)A in the Transcriptome: m(6)A-Binding Proteins. Trends Cell Biol, 28, 113–127.

Petersen, G., Cuenca, A., Moller, I.M. and Seberg, O. (2015) Massive gene loss in mistletoe (Viscum, Viscaceae) mitochondria. Scientific reports, 5, 1–7.

Quinlan, A.R. and Hall, I.M. (2010) BEDTools: a flexible suite of utilities for comparing genomic features. Bioinformatics, 26, 841–842.

Růžička, K., Zhang, M., Campilho, A., Bodi, Z., Kashif, M., Saleh, M., Eeckhout, D., El-Showk, S., Li, H., Zhong, S., De Jaeger, G., Mongan, N.P., Hejatko, J., Helariutta, Y. and Fray, R.G. (2017) Identification of factors required for m(6) A mRNA methylation in Arabidopsis reveals a role for the conserved E3 ubiquitin ligase HAKAI. New Phytol, 215, 157–172.

Sabar, M., Gagliardi, D., Balk, J. and Leaver, C.J. (2003) ORFB is a subunit of F1F(O)-ATP synthase: insight into the basis of cytoplasmic male sterility in sunflower. EMBO Rep, 4, 381–386.

Schallenberg-Rüdinger, M. and Knoop, V. (2016) Coevolution of Organelle RNA Editing and Nuclear Specificity Factors in Early Land Plants. In Advances in Botanical Research (Rensing, S.A. ed: Academic Press, pp. 37–93.

Schertl, P. and Braun, H.P. (2014) Respiratory electron transfer pathways in plant mitochondria. Front Plant Sci, 5, 163.

Senkler, J., Rugen, N., Eubel, H., Hegermann, J. and Braun, H.-P. (2018) Absence of Complex I Implicates Rearrangement of the Respiratory Chain in European Mistletoe. Current Biology, 28, 1606–1613.e1604.

Shen, L., Liang, Z., Gu, X., Chen, Y., Teo, Z.W., Hou, X., Cai, W.M., Dedon, P.C., Liu, L. and Yu, H. (2016) N6-Methyladenosine RNA Modification Regulates Shoot Stem Cell Fate in Arabidopsis. Dev Cell.

Shikanai, T. (2015) RNA editing in plants: Machinery and flexibility of site recognition. Biochim Biophys Acta, 1847, 779–785.

Skippington, E., Barkman, T.J., Rice, D.W. and Palmer, J.D. (2015) Miniaturized mitogenome of the parasitic plant Viscum scurruloideum is extremely divergent and dynamic and has lost all nad genes. Proceedings of the National Academy of Sciences, 112, E3515–E3524.

Sloan, D., Alverson, A., Chuckalovcak, J., Wu, M., McCauley, D., Palmer, J. and Taylor, D. (2012) Rapid evolution of enormous, multichromosomal genomes in flowering plant mitochondria with exceptionally high mutation rates. PLoS Biol, 10, e1001241.

Sloan, D.B., Wu, Z. and Sharbrough, J. (2018) An improved mitochondrial reference genome for Arabidopsis thaliana Col-0. bioRxiv, 249086.

Slobodin, B., Han, R., Calderone, V., Vrielink, J.A., Loayza-Puch, F., Elkon, R. and Agami, R. (2017) Transcription Impacts the Efficiency of mRNA Translation via Co-transcriptional N6-adenosine Methylation. Cell, 169, 326–337.e312.

Small, I. (2013) Mitochondrial genomes as living ’fossils’. BMC Biology, 11, 30.

Sultan, L.D., Mileshina, D., Grewe, F., Rolle, K., Abudraham, S., Głodowicz, P., Khan Niazi, A., keren, I., Shevtsov, S., Klipcan, L., Barciszewski, J., Mower, J.P., Dietrich, A. and Ostersetzer, O. (2016) The reverse-transcriptase/RNA-maturase protein MatR is required for the splicing of various group II introns in Brassicaceae mitochondria. Plant Cell, 28, 2805–2829.

Sun, H., Zhang, M., Li, K., Bai, D. and Yi, C. (2019) Cap-specific, terminal N6-methylation by a mammalian m6Am methyltransferase. Cell research, 29, 80–82.

Takenaka, M., Verbitskiy, D., van der Merwe, J.A., Zehrmann, A. and Brennicke, A. (2008) The process of RNA editing in plant mitochondria. Mitochondrion, 8, 35–46.

Thuring, K., Schmid, K., Keller, P. and Helm, M. (2016) Analysis of RNA modifications by liquid chromatography-tandem mass spectrometry. Methods, 107, 48–56.

Unseld, M., Marienfeld, J.R., Brandt, P. and Brennicke, A. (1997) The mitochondrial genome of Arabidopsis thaliana contains 57 genes in 366,924 nucleotides. Nature Genet, 15, 57–61.

Uyttewaal, M., Mireau, H., Rurek, M., Hammani, K., Arnal, N., Quadrado, M. and Giegé, P. (2008) PPR336 is associated with polysomes in plant mitochondria. Journal of Molecular Biology, 375, 626–636.

Visvanathan, A. and Somasundaram, K. (2018) mRNA Traffic Control Reviewed: N6-Methyladenosine (m(6) A) Takes the Driver’s Seat. Bioessays, 40.

Wahleithner, J.A., MacFarlane, J.L. and Wolstenholme, D.R. (1990) A sequence encoding a maturase-related protein in a group II intron of a plant mitochondrial nad1 gene. PNAS, 87, 548–552.

Waltz, F., Nguyen, T.-T., Arrivé, M., Bochler, A., Chicher, J., Hammann, P., Kuhn, L., Quadrado, M., Mireau, H., Hashem, Y. and Giegé, P. (2019) Small is big in Arabidopsis mitochondrial ribosome. Nature Plants, 5, 106–117.

Wan, Y., Tang, K., Zhang, D., Xie, S., Zhu, X., Wang, Z. and Lang, Z. (2015) Transcriptome-wide high-throughput deep m6A-seq reveals unique differential m6A methylation patterns between three organs in Arabidopsis thaliana. Genome Biol, 16, 272.

Wang, X., Lu, Z., Gomez, A., Hon, G.C., Yue, Y., Han, D., Fu, Y., Parisien, M., Dai, Q., Jia, G., Ren, B., Pan, T. and He, C. (2014) N6-methyladenosine-dependent regulation of messenger RNA stability. Nature, 505, 117–120.

Wang, X., Ryu, D., Houtkooper, R.H. and Auwerx, J. (2015a) Antibiotic use and abuse: A threat to mitochondria and chloroplasts with impact on research, health, and environment. Bioessays, 37, 1045–1053.

Wang, X., Zhao, B.S., Roundtree, I.A., Lu, Z., Han, D., Ma, H., Weng, X., Chen, K., Shi, H. and He, C. (2015b) N(6)-methyladenosine Modulates Messenger RNA Translation Efficiency. Cell, 161, 1388–1399.

Wang, Z., Tang, K., Zhang, D., Wan, Y., Wen, Y., Lu, Q. and Wang, L. (2017) High-throughput m6A-seq reveals RNA m6A methylation patterns in the chloroplast and mitochondria transcriptomes of Arabidopsis thaliana. PLoS One, 12, e0185612.

Woodson, J.D. and Chory, J. (2008) Coordination of gene expression between organellar and nuclear genomes. Nat Rev Genet, 9, 383–395.

Xiao, W., Adhikari, S., Dahal, U., Chen, Y.S., Hao, Y.J., Sun, B.F., Sun, H.Y., Li, A., Ping, X.L., Lai, W.Y., Wang, X., Ma, H.L., Huang, C.M., Yang, Y., Huang, N., Jiang, G.B., Wang, H.L., Zhou, Q., Wang, X.J., Zhao, Y.L. and Yang, Y.G. (2016) Nuclear m(6)A Reader YTHDC1 Regulates mRNA Splicing. Mol Cell, 61, 507–519.

Yue, Y., Liu, J. and He, C. (2015) RNA N6-methyladenosine methylation in post-transcriptional gene expression regulation. Genes Dev, 29, 1343–1355.

Zhong, S., Li, H., Bodi, Z., Button, J., Vespa, L., Herzog, M. and Fray, R.G. (2008) MTA Is an Arabidopsis Messenger RNA Adenosine Methylase and Interacts with a Homolog of a Sex-Specific Splicing Factor. The Plant Cell, 20, 1278–1288.

Zmudjak, M. and Ostersetzer-Biran, O. (2017) RNA METABOLISM AND TRANSCRIPT REGULATION. In Annual Plant Reviews, Volume 50 pp. 261–309, D. C. Logan (Ed.).

Zmudjak, M., Shevtsov, S., Sultan, L.D., Keren, I. and Ostersetzer-Biran, O. (2017) Analysis of the Roles of the Arabidopsis nMAT2 and PMH2 Proteins Provided with New Insights into the Regulation of Group II Intron Splicing in Land-Plant Mitochondria. International journal of molecular sciences, 18, 1–25.

Zoschke, R., Watkins, K.P. and Barkan, A. (2013) A rapid ribosome profiling method elucidates chloroplast ribosome behavior in vivo. The Plant cell, 25, 2265–2275.

